# Goal-specific brain MRI harmonization

**DOI:** 10.1101/2022.03.05.483077

**Authors:** Lijun An, Jianzhong Chen, Pansheng Chen, Chen Zhang, Tong He, Christopher Chen, Juan Helen Zhou, B.T. Thomas Yeo, the Alzheimer’s Disease Neuroimaging Initiative, the Australian Imaging Biomarkers and Lifestyle Study of Aging

## Abstract

There is significant interest in pooling magnetic resonance image (MRI) data from multiple datasets to enable mega-analysis. Harmonization is typically performed to reduce heterogeneity when pooling MRI data across datasets. Most MRI harmonization algorithms do not explicitly consider downstream application performance during harmonization. However, the choice of downstream application might influence what might be considered as study-specific confounds. Therefore, ignoring downstream applications during harmonization might potentially limit downstream performance. Here we propose a goal-specific harmonization framework that utilizes downstream application performance to regularize the harmonization procedure. Our framework can be integrated with a wide variety of harmonization models based on deep neural networks, such as the recently proposed conditional variational autoencoder (cVAE) harmonization model. Three datasets from three different continents with a total of 2787 participants and 10085 anatomical T1 scans were used for evaluation. We found that cVAE removed more dataset differences than the widely used ComBat model, but at the expense of removing desirable biological information as measured by downstream prediction of mini mental state examination (MMSE) scores and clinical diagnoses. On the other hand, our goal-specific cVAE (gcVAE) was able to remove as much dataset differences as cVAE, while improving downstream cross-sectional prediction of MMSE scores and clinical diagnoses.

## 1 Introduction

Large scale MRI datasets from multiple sites have boosted the study of human brain structure and function (Yeo et al., 2011; Van Essen et al., 2013; Miller et al., 2016; Volkow et al., 2018). Combining datasets from multiple sites can potentially boost statistical power, so there is significant interest in pooling data across multiple sites (Thompson et al., 2017; Whelan et al., 2018; Tang et al., 2020; Lu et al., 2020). However, MRI data is sensitive to variation of scanners across different sites (Jovicich et al., 2006; Magnotta et al., 2012; Chen et al., 2014; Hawco et al., 2018), so post-acquisition harmonization is necessary for removing unwanted variabilities in pooling data across multiple studies.

A popular harmonization approach is the ComBat framework (Fortin et al., 2017, 2018; Yu et al., 2018) that utilizes a mixed effects regression model to remove additive and multiplicative site effects. Other ComBat variants have since been proposed (Garcia-Dias et al., 2020; Pomponio et al., 2020; Wachinger et al., 2021). However, most ComBat variants consider each brain region separately (but see Chen et al., 2019), so might not be able to remove nonlinear site differences that are distributed across brain regions.

These nonlinear distributed site differences might be more readily removed by harmonization approaches based on deep neural networks (DNNs; (Tanno et al., 2017; Ning et al., 2019; Blumberg et al., 2019). One popular approach is the use of the variational autoencoder (VAE) framework (Moyer et al., 2020; Russkikh et al., 2020; Zuo et al., 2021), which typically uses an encoder to generate site-invariant latent representations. Site information can then be added to the latent representations to “reconstruct” the MRI data. Another popular approach is the use of generative adversarial networks and cycle consistency constraints (Zhu et al., 2017; Zhao et al., 2019; Dewey et al., 2019; Modanwal et al., 2020; Bashyam et al., 2021).

However, most previously proposed harmonization approaches do not consider downstream applications in the harmonization procedure. It is important to note that the goal of MRI harmonization is to remove ‘unwanted’ dataset differences, while preserving relevant biological information. However, unwanted dataset differences depend on the application. For example, if our goal is to develop an Alzheimer’s disease (AD) dementia prediction model that is generalizable across different racial groups, then ‘race’ might be considered an undesirable study difference. On the other hand, if we are interested in studying AD progression across different racial groups, then racial information needs to be preserved in the harmonization process. Therefore, ignoring downstream applications in the harmonization procedure might potentially limit downstream performance.

In this study, we propose a goal-specific harmonization framework that utilizes downstream applications to regularize the harmonization model. Our approach can be integrated with most DNN-based harmonization approaches, such as the conditional VAE (cVAE) harmonization model (Moyer et al., 2020), which was previously applied to diffusion MRI data. We then compared the resulting goal-specific cVAE (gcVAE) model with cVAE and ComBat using three datasets comprising 2787 participants and 10085 anatomical MRI scans. The evaluation procedure tested the ability of different harmonization models to remove dataset differences while retaining biological information as measured by downstream cross-sectional prediction of mini mental state examination (MMSE) scores and clinical diagnoses.

### 2.1 Datasets

## 2 Methods

In this study, we considered T1 structural MRI data from the Alzheimer’s Disease Neuroimaging Initiative (ADNI) (Jack et al., 2008, 2010), the Australian Imaging, Biomarkers and Lifestyle (AIBL) study (Ellis et al., 2009, 2010) and the Singapore Memory Ageing and Cognition Centre (MACC) Harmonization cohort (Hilal et al., 2015; Chong et al., 2017; Hilal et al., 2020). Data collection was approved by the Institutional Review Board (IRB) at each corresponding institution. The analysis in the current study is approved by the National University of Singapore IRB. Across all three datasets, MRI data was collected at multiple timepoints.

In the case of ADNI (Jack et al., 2008, 2010), we considered data from ADNI1 and ADNI2/Go. For ADNI1, the MRI scans were collected from 1.5 and 3T scanners from different vendors (see Table S1 for more details). For ADNI2/Go, the MRI scans were collected from 3T scanners. There were 1735 participants with at least one T1 MRI scan. There was a total of 7955 MRI scans across the different timepoints of the 1735 participants.

In the case of AIBL (Ellis et al., 2009, 2010), the MRI scans were collected from 1.5T and 3T Siemens (Avanto, Tim Trio and Verio) scanners (see Table S2 for more details).There were 495 participants with at least one T1 MRI scan. There was a total of 933 MRI scans across the different timepoints of the 495 participants.

In the case of MACC (Hilal et al., 2015; Chong et al., 2017; Hilal et al., 2020), the MRI scans were collected from a Siemens 3T Tim Trio scanner. There were 557 participants with at least one T1 MRI scan. There was a total of 1197 MRI scans across the different timepoints of the 557 participants.

### 2.2 Data Preprocessing

Our goal is to harmonize volumes of regions of interest (ROIs) across datasets. Here, 108 cortical and subcortical ROIs were defined based on the FreeSurfer software (Fischl et al., 2002; Desikan et al., 2006). In the case of ADNI, we utilized the ROI volumes provided by ADNI. These ROIs were generated by ADNI after several preprocessing steps (http://adni.loni.usc.edu/methods/mri-tool/mri-pre-processing/) followed by the FreeSurfer version 4.3 (ADNI1) and 5.1 (ADNI2/GO) recon-all pipeline. In the case of AIBL and MACC, FreeSurfer version 6.0 recon-all pipeline was utilized. Therefore, differences between the datasets arose from both scanner and preprocessing differences.

### 2.3 Workflow overview

In this study, we sought to harmonize brain ROI volumes between ADNI and AIBL, as well as ADNI and MACC. Figures 1 and 2 illustrate the workflow in this study using AIBL as an illustration. The procedure is exactly the same for MACC.

**Figure 1.**
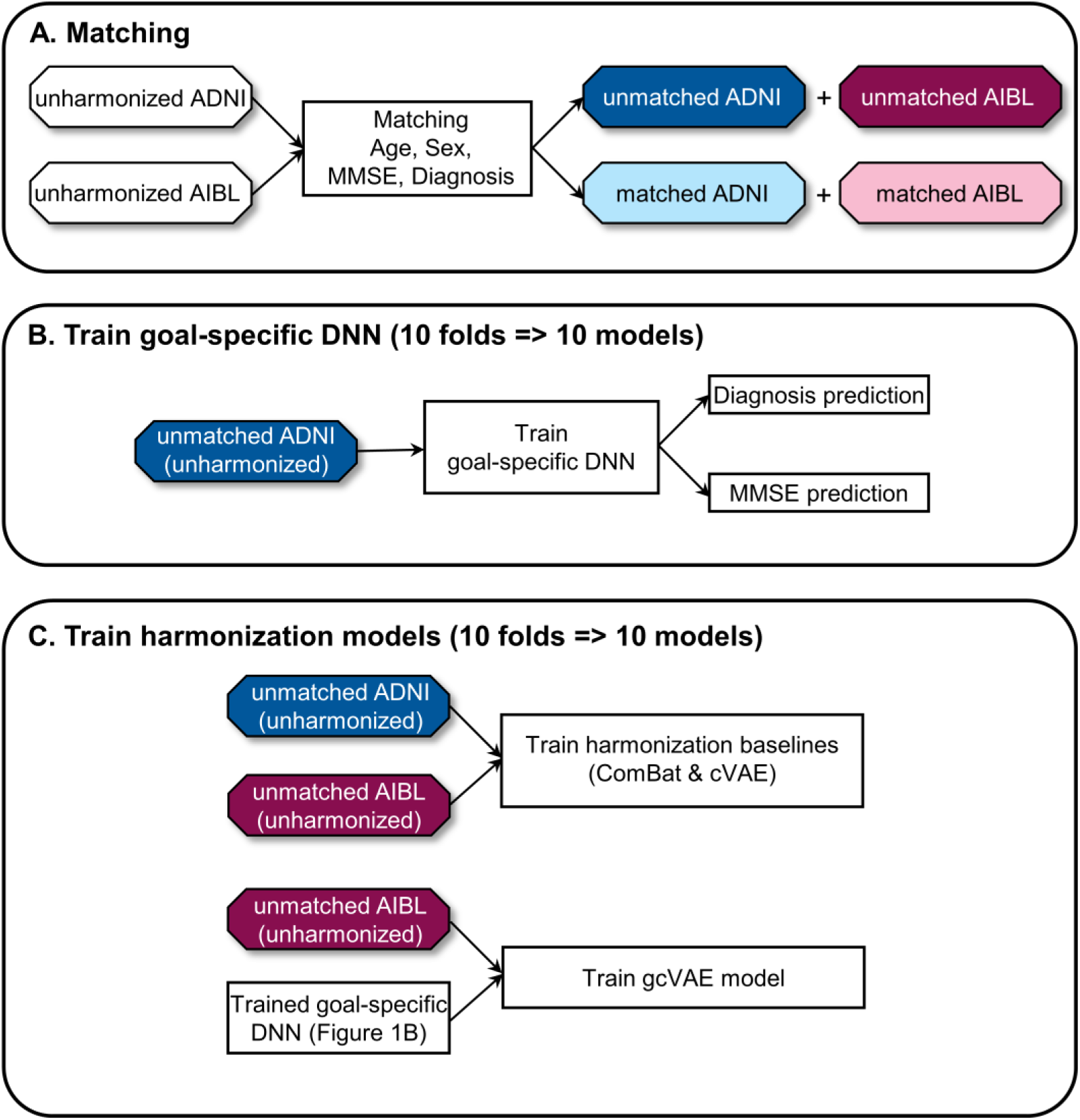
Workflow of current study for data matching and model training. We illustrate the workflow using ADNI and AIBL. The same procedure was applied to ADNI and MACC. (A) Matching participants to derive test set for harmonization evaluation. (B) Train goal-specific deep neural network (DNN) using unmatched unharmonized ADNI data to predict clinical diagnosis and MMSE. (C) Train harmonization models using unmatched unharmonized data. We note that ComBat and cVAE were trained using unmatched unharmonized ADNI and AIBL data, while gcVAE was trained using unmatched unharmonized AIBL data and the goal-specific DNN (from Figure 1B). Dark colors (e.g., dark red and dark blue) are used to indicate unmatched participants, while light colors (e.g., pink and light blue) are used to indicate matched participants.

**Figure 2.**
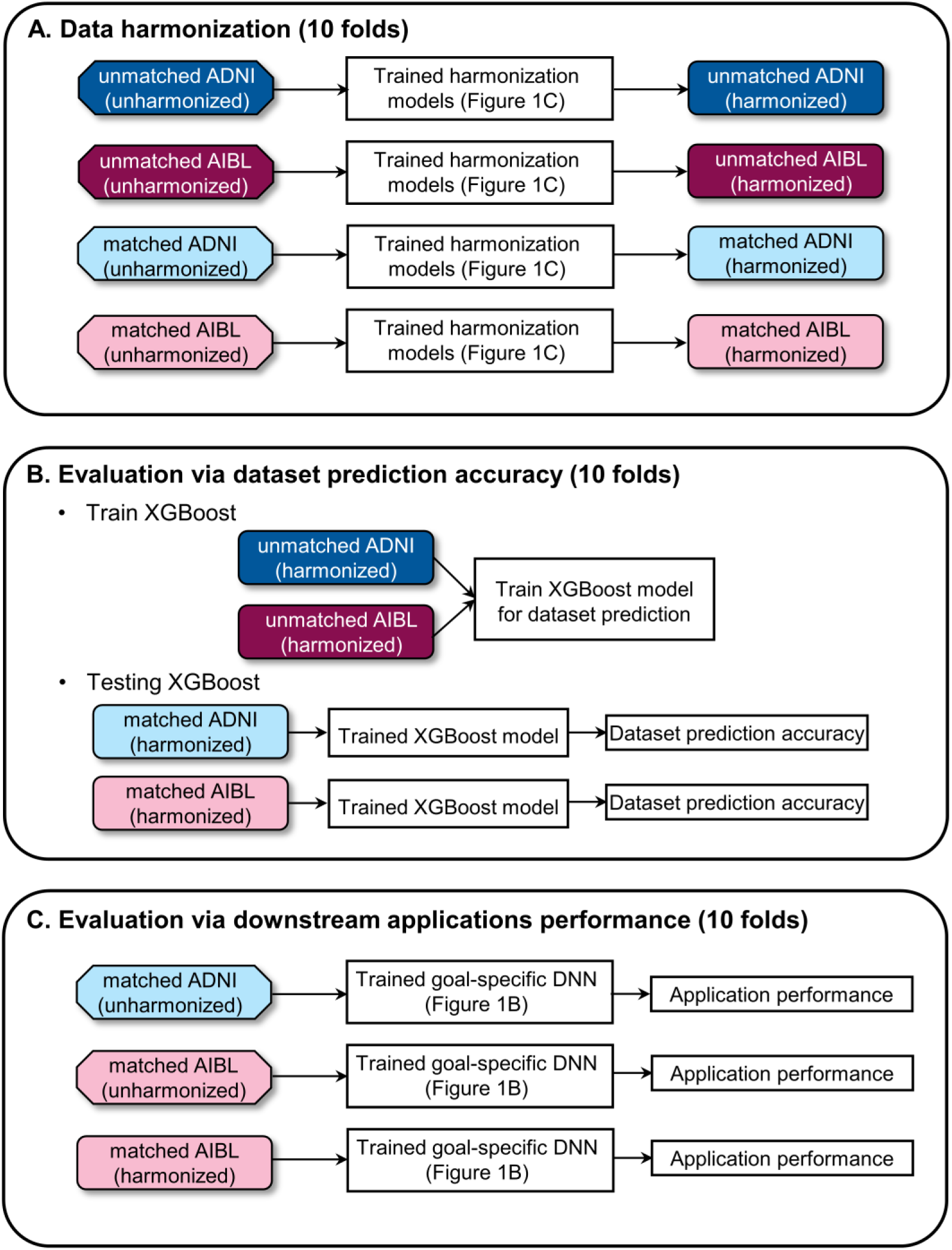
Workflow of current study for data harmonization and performance evaluation. We illustrate the workflow using ADNI and AIBL. The same procedure was applied to ADNI and MACC. (A) Harmonize data using trained harmonization models from Figure 1C. (B) Evaluate harmonization performance using XGBoost dataset prediction model. (C) Evaluate harmonization performance using goal-specific DNN (Figure 1B) to predict MMSE and clinical diagnosis. We note that dark colors (e.g., dark red and dark blue) are used to indicate unmatched participants, while light colors (e.g., pink and light blue) are used to indicate matched participants. On the other hand, octagons are used to indicate unharmonized data, while rectangles (with rounded corners) are used to indicate harmonized data.

In the case of AIBL, we used the Hungarian matching algorithm (Kuhn, 1955) to first select pairs of ADNI and AIBI participants with matched number of timepoints, age, sex, MMSE and clinical diagnosis (Figure 1A). The distributions of age, sex, MMSE and clinical diagnosis of all participants and matched participants are shown in Figure 3.

**Figure 3.**
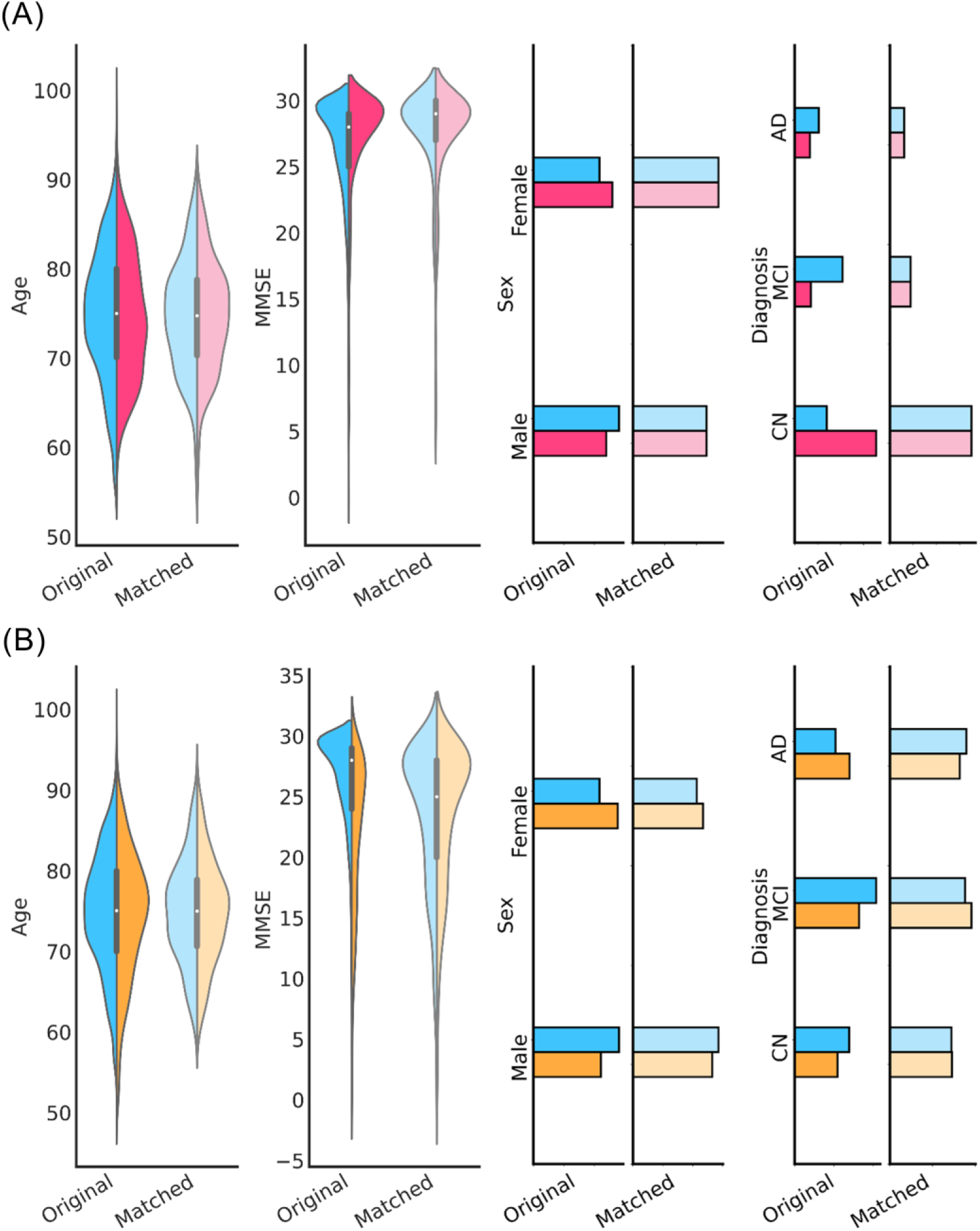
Age, MMSE, sex and clinical diagnosis distributions before and after matching. (A) Distributions of age, sex, MMSE and clinical diagnosis for ADNI (blue) and AIBL (red). Differences in the attributes between ADNI and AIBL were not significant after matching. (B) Distributions of age, sex, MMSE and clinical diagnosis for ADNI (blue) and MACC (yellow). Differences in the attributes between ADNI and MACC were not significant after matching. P values showing the quality of the matching procedure are found in Tables S3 to S9.

There were 247 pairs of matched AIBI and ADNI participants with an average of 1.1 scans per participant. The same approach was applied to ADNI and MACC, yielding 277 pairs of matched MACC and ADNI participants with an average of 1.5 scans per participant. We note that not all timepoints have corresponding MMSE and clinical diagnosis information. Therefore, care was taken to ensure that all timepoints in the matched participants had both MMSE and clinical diagnosis. Care was taken to ensure that all scans of every participant were classified as either “matched” or “unmatched”, and not split between the two categories. P values showing the quality of the matching procedure are found in Tables S3 to S9.

The unmatched ADNI data was used to train goal-specific deep neural networks (DNN) for predicting MMSE and clinical diagnosis (Figure 1B; details in Section 2.5). Here, clinical diagnosis categories were normal controls, mild cognitive impairment, and Alzheimer’s disease dementia. The clinical diagnoses from all three datasets were determined by multiple criteria, including MRI and cognitive tests. The unmatched ADNI and AIBL participants were also used to fit ComBat and cVAE (Figure 1C; details in Section 2.6). The unmatched AIBL participants and goal-specific DNN (from Figure 1B) were utilized for training the gcVAE model (Figure 1C). The same procedure was applied to ADNI and MACC.

The trained harmonization models were then applied to unharmonized brain volumes of all matched and unmatched participants (Figure 2A). The harmonized data was evaluated with two criteria (Figures 2B and 2C). The first criterion was dataset prediction performance, in which a machine learning algorithm was used to predict which dataset the harmonized data came from (Figure 2B). Lower dataset prediction performance indicates better harmonization. More specifically, we trained a XGBoost classifier (Chen & Guestrin, 2016) using the harmonized ADNI and harmonized AIBL brain volumes from the unmatched participants (Figure 2B). We then applied the classifier to the harmonized ADNI and AIBL brain volumes from the matched participants (details in Section 2.8). The same procedure was applied to ADNI and MACC.

However, a simple way to achieve perfect dataset prediction results was to map all brain volumes to zero, thus losing all biological information. Therefore, the second criterion was downstream application performance (Figure 2C). Here, we applied the goal-specific DNN (from Figure 1B) to the harmonized AIBL brain volumes from the matched participants. To demonstrate the effects of no harmonization, the goal-specific DNN was also applied to the unharmonized AIBL and unharmonized ADNI brain volumes from the matched participants (Figure 2C). The same procedure was applied to ADNI and MACC.

We note that the goal-specific DNN (Figure 1B), harmonization models (Figure 1C) and dataset prediction classifier (Figure 2B) were trained on unmatched data, while harmonization evaluation was performed on matched data (Figures 2B and 2C). The matching procedure was important to ensure that prediction performance was comparable between matched ADNI and matched AIBI participants. Suppose we did the opposite: trained a clinical diagnosis prediction model on matched ADNI participants and then tested the model on unmatched ADNI and unmatched AIBL participants. In this scenario, the clinical diagnosis prediction performance would not be comparable between unmatched ADNI and unmatched AIBL participants. More specifically, suppose unmatched ADNI comprised mostly participants with AD and healthy participants, as well as few participants with mild cognitive impairment (MCI). On the other hand, suppose AIBL contained equal proportions of healthy participants, participants with MCI and participants with AD. In this scenario, because it is easier to distinguish between healthy controls and participants with AD, compared with distinguishing participants with MCI from the other two classes (participants with AD and healthy participants), the prediction performance would likely be better in unmatched ADNI compared with unmatched AIBL, even if there was no scanner difference between the two sites. By testing prediction performance on matched AIBL and matched ADNI participants, we ensure that any drop in prediction performance was due to unavoidable site differences, such as scanner differences.

### 2.4 Training, validation and test procedure

As mentioned in the previous section, the matched participants were used as the test set for evaluation (Figure 2C). The unmatched participants were used for training the goal-specific DNN (Figure 1B), harmonization (Figure 1C) and dataset prediction (Figure 2B) models. More specifically, we divided the unmatched participants into 10 groups. Recall that a participant might be scanned at multiple timepoints. Care was taken to ensure that all timepoints of any participant were assigned to be in a single group, and not split across multiple groups.

To train the goal-specific DNN, harmonization and dataset prediction models, 9 groups were used for training, while the remaining group was used as a validation set to tune the hyperparameters. This procedure was repeated 10 times with a different group being the validation set. Therefore, we ended up with 10 sets of trained models. The 10 sets of harmonization models were applied to the unharmonized data (Figure 2A), yielding 10 sets of harmonized data. The 10 sets of XGBoost classifiers and goal-specific DNNs were applied to the 10 corresponding sets of harmonized data for evaluation (Figures 2B and 2C).

### 2.5 Goal-specific DNNs

Here we utilized DNNs to predict MMSE and clinical diagnosis (normal controls, mild cognitive impairment or Alzheimer’s disease dementia) jointly. The goal-specific DNNs were used to train the gcVAE model (Figure 1C) and evaluate the harmonization approaches (Figure 2C). The inputs to the goal-specific DNNs were the brain ROI volumes. 10 DNNs were trained with a 10-fold cross-validation procedure (Section 2.4) using the *unmatched unharmonized* ADNI MRI volumes (Figure 1B). The training procedure utilized the unharmonized ADNI data without differentiation among ADNI sites.

Recall that not all unmatched timepoints had MMSE and clinical diagnosis information. Therefore, we used the previous timepoint with available information to fill in the missing data (Lipton et al., 2016; Che et al., 2018; Nguyen et al., 2020). Note that this filling in procedure was only performed during training procedure for the unmatched participants.

The architecture of the goal-specific DNN was a generic feedforward neural network, where every layer was fully connected with the next layer. The nonlinear activation function ReLU (Maas et al., 2013) was utilized. The DNN loss function corresponded to the weighted sum of the mean absolute error (MAE) for MMSE prediction and cross entropy loss for clinical diagnosis prediction: L_goalDNN_ = λ_MMSE_ MAE + λ_DX_ CrossEntropy. λ_MMSE_ and λ_DX_ were two hyperparameters that were tuned on the validation set.

The metric for tuning hyperparameters in the validation set was the weighted sum of MMSE MAE and clinical diagnosis accuracy: ½ MAE – Diagnosis Accuracy. The MAE term was divided by two so the two terms had similar ranges of values. We utilized the HORD algorithm (Regis & Shoemaker, 2013; Ilievski et al., 2017; Eriksson et al., 2020) to find the best set of hyperparameters using the validation set (Table 1). The trained DNN after 100 epochs was utilized for subsequent analyses.

**Table 1.** Hyperparameters estimated from the validation set. We note that a learning rate decay strategy was utilized. After K training epochs (where K = learning rate step), the learning rate was reduced by a factor of 10.

| Hyperparameter | Search range |
| --- | --- |
| Initial learning rate | 1e-4 – 1e-3 |
| Learning rate step | 10 – 99 |
| Dropout rate | 0 – 0.5 |
| $\lambda_{MMSE}$ | 0 – 1 |
| $\lambda_{DX}$ | 0 – 1 |
| Nodes for each layer | 32 – 512 |
| Number of layers | 2 – 5 |

At the evaluation phase (Figure 2C), the 10 goal-specific DNNs were applied to the harmonized brain volumes from the matched AIBL participants, as well as unharmonized brain volumes from the matched AIBL and ADNI participants. The prediction performance was averaged across all time points of each participant and the 10 goal-specific DNNs before averaging across participants. The same procedure was applied to ADNI and MACC participants.

### 2.6 Baseline harmonization models

Here, we considered ComBat (Johnson et al., 2007) and cVAE (Moyer et al., 2020) as baseline models.

#### 2.6.1 ComBat

ComBat is a linear mixed effects model that controls for additive and multiplicative site effects (Johnson et al., 2007). Here we utilized the R implementation of the algorithm (https://github.com/Jfortin1/ComBatHarmonization). The ComBat model is as follows:

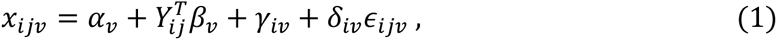

where *i* is the site index, *j* is the participant index and *v* is the brain ROI index. *x*_*ijv*_ is the volume of the *v*-th brain ROI of subject *j* from site *i*. *γ*_*iv*_ is the addictive site effect. *δ*_*iv*_ is the multiplicative site effect. *ε*_*ijv*_ is the residual error term following a normal distribution with zero mean and variance 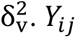 are the covariates of subject *j* from site *i*.

The ComBat parameters *α*_*v*_, *β*_*v*_, *γ*_*iv*_ and *δ*_*iv*_ were estimated for each brain ROI using the unmatched unharmonized ROI volumes (Figure 1C). The estimated parameters can then be applied to a new participant *i* from site *j* with brain volume *x*_*ijv*_ and covariates *Y*_*ij*_

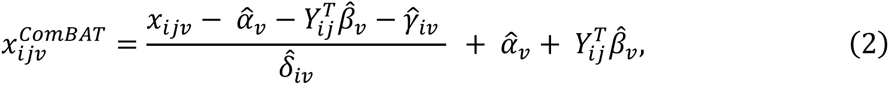

where ^ indicates that the parameter was estimated from the *unmatched unharmonized* ROI volumes from ADNI and AIBL. A separate ComBat model was fitted for ADNI and MACC brain volumes. Observe that the equation required the covariates of the new participant. Given that we would like to predict MMSE and clinical diagnosis in the matched participants, this implied that MMSE and clinical diagnosis information were not available in the matched participants. Therefore, we could not utilize MMSE and clinical diagnosis as covariates in the ComBat model. Therefore, in the main results, we only utilized age and sex as covariates. However, as a control analysis (Section 2.9.3), we also considered a version of ComBat where age, sex, MMSE and clinical diagnoses were used as covariates.

Furthermore, since the goal-specific DNNs were trained with unmatched unharmonized ADNI data without distinguishing among the sites (Section 2.5), for consistency, the ComBat procedure also treated ADNI as a single site despite the data coming from multiple sites and scanners. This was also the case for AIBL.

Note that equation (2) mapped both ADNI and AIBL data to an “intermediate” space, which is not an issue for the purpose of dataset prediction because the XGBoost classifier was trained from scratch (Figure 2B; Section 2.8). However, for the purpose of predicting MMSE and clinical diagnosis, since the goal-specific DNN was trained with unharmonized ADNI data, we used the ref.batch option in the ComBat package to map AIBL data to “ADNI-space” after harmonization. The same procedure was applied to ADNI and MACC.

#### 2.6.2 cVAE

The conditional variational autoencoder (cVAE) model was proposed by Moyer and colleagues to harmonize diffusion MRI data (Moyer et al., 2020). Here, we applied cVAE to harmonize brain ROI volumes. The cVAE model is illustrated in Figure 4A. Input brain volumes were passed through an encoder DNN yielding representation *z*. Site index *s* was concatenated with the latent representation *z* before feeding into the decoder DNN, resulting in the reconstructed brain volumes 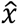. By incorporating the mutual information *I*(*z*, *s*) in the cost function, this encouraged the learned representation *z* to be independent of the site *s*.

**Figure 4.**
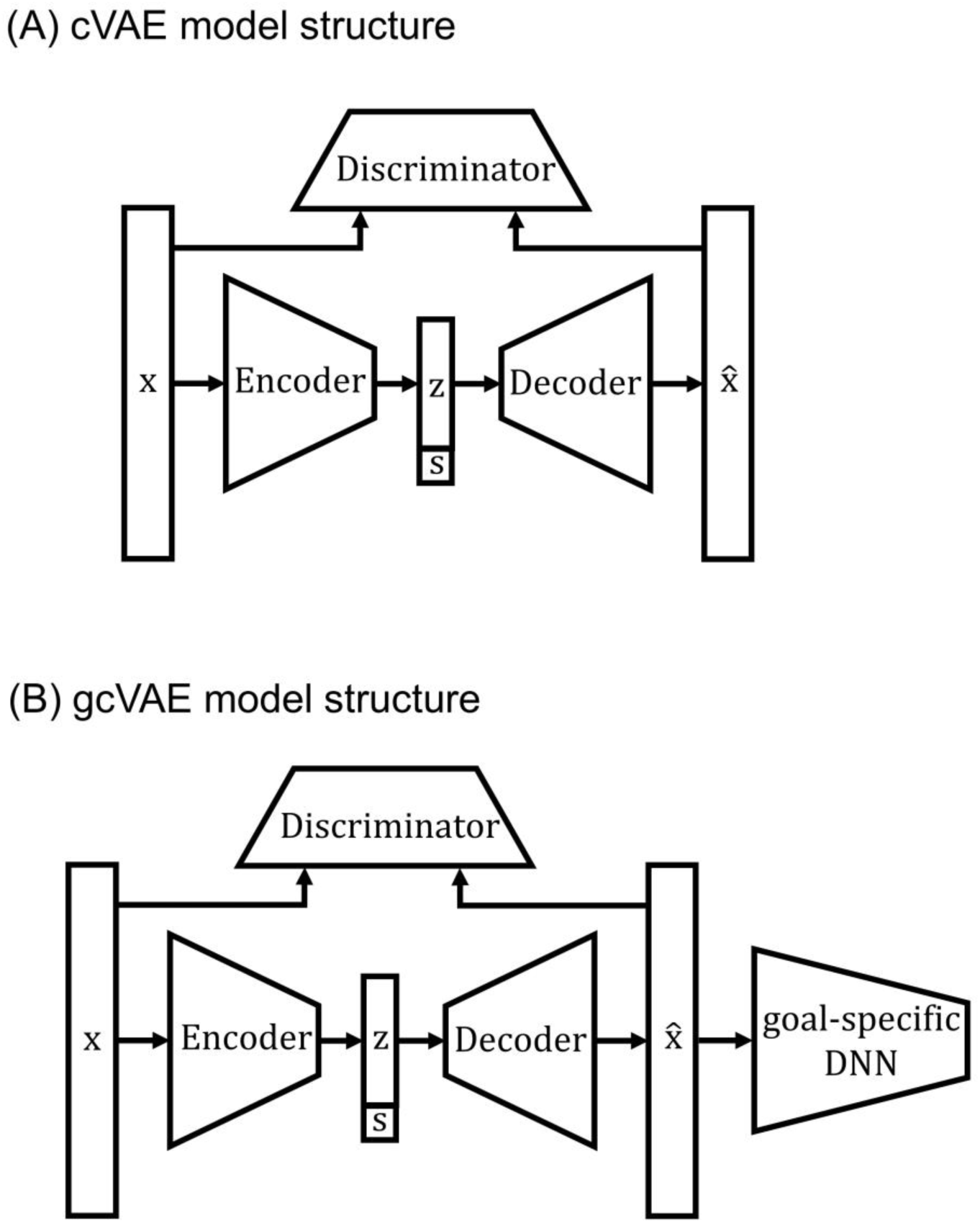
cVAE and gcVAE model structures. **(A)** Model structure for the cVAE model. Encoder, decoder, and discriminator were all fully connected feedforward DNNs. *s* was the site we wanted to map the brain volumes to. (B) Model structure for the gcVAE model. The goal-specific DNN from Section 2.5 (Figure 1B) was used to guide the cVAE harmonization process. During training of gcVAE, the weights of the goal-specific DNN were fixed.

The resulting lost function is as follows:

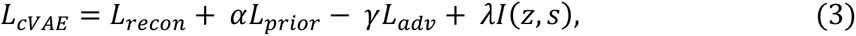

where *L*_*recon*_ is the mean square error (MSE) between *x* and 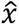, so this encouraged the harmonized volumes to be similar to the unharmonized volumes. To further encourage *x* and 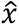 to be similar, Moyer and colleagues added an additional term *L*_*adv*_, which is the soft-max cross-entropy loss of an adversarial discriminator seeking to distinguish between *x* and 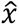. Finally, *L*_*prior*_ is the standard KL divergence between representation *z* and the multivariate Gaussian distribution with zero mean and identity covariance matrix (Sohn et al., 2015).

Both the decoder and encoder were instantiated as generic feedforward neural networks, where every layer was fully connected with the next layer. Following Moyer and colleagues, the nonlinear activation function tanh (Maas et al., 2013) was utilized. During the training process, *s* is the true site information for input brain volumes *x*. After training, we could map input *x* to any site by changing *s*. The metric for tuning hyperparameters in the validation set was the weighted sum of the reconstruction loss (MSE between *x* and 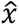) and the subject-level dataset prediction accuracy: ½ MAE + Dataset Accuracy. The MAE reconstruction loss was divided by two so the two terms had similar ranges of values. Dataset prediction accuracy was obtained by training a XGBoost classifier on the training set and applying to the validation set. We utilized the HORD algorithm (Regis & Shoemaker, 2013; Ilievski et al., 2017; Eriksson et al., 2020) to find the best set of hyperparameters using the validation set (Table 2). The trained DNN after 1000 epochs was utilized for subsequent analyses.

**Table 2.** Hyperparameters estimated from the validation set. We note that a learning rate decay strategy was utilized. After K training epochs (where K = learning rate step), the learning rate was reduced by a factor of 10.

| Hyperparameter | Search range |
| --- | --- |
| Initial learning rate | 1e-2 – 1e-1 |
| Learning rate step | 10 - 999 |
| Dropout rate | 0 – 0.5 |
| $\alpha$ | 0.01 - 1 |
| $\gamma$ | 0.01 - 10 |
| $\lambda$ | 0.01 - 1 |
| Nodes for each layer | 32 - 512 |
| Number of layers | 2 - 4 |
| Node for $z$ | 32 - 512 |

Similar to ComBat, the cVAE model was trained using *unmatched unharmonized* brain volumes from ADNI and AIBL. A separate model was trained using ADNI and MACC. For consistency, the cVAE model also treated ADNI and AIBL as single sites.

Similar to ComBat, for the purpose of dataset prediction, data were mapped to intermediate space by setting the site *s* to 0 during harmonization. On the other hand, for the purpose of predicting MMSE and clinical diagnosis, data from AIBL (and MACC) was mapped to ADNI space by setting the site *s* to correspond to ADNI.

### 2.7 Goal-specific cVAE (gcVAE)

To incorporate downstream application performance in the harmonization procedure, the outputs of the cVAE (Figure 4A) were fed into the goal-specific DNN (Section 2.5). The resulting goal-specific cVAE (gcVAE) is illustrated in Figure 4B. The loss function of the gcVAE was given by corresponded to the weighted sum of the mean absolute error (MAE) for MMSE prediction and cross entropy loss for clinical diagnosis prediction:

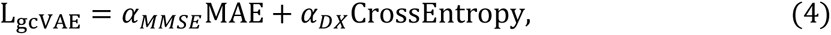

where *α*_*MMSE*_ and *α*_*DX*_ were two hyperparameters to be tuned with the validation set. The loss function was used to finetune the trained cVAE model (Section 2.6.2) using the training set with a relatively small learning rate. We note that the weights of the goal-specific DNN model were frozen during this finetuning procedure.

The metric for tuning hyperparameters in the validation set was the weighted sum of MMSE MAE and clinical diagnosis accuracy: ½ MAE – Diagnosis Accuracy (same as Section 2.5). Since there were only three hyperparameters (learning rate, *α*_*MMSE*_ and *α*_*DX*_), a grid search was performed using the validation set to find the best set of hyperparameters.

The gcVAE model was trained using *unmatched unharmonized* brain volumes from AIBL. A separate model was trained using ADNI and MACC. For consistency, the gcVAE model also treated ADNI and AIBL as single sites.

Similar to ComBat, for the purpose of dataset prediction, data were mapped to intermediate space by setting the site *s* to 0 during harmonization. On the other hand, for the purpose of predicting MMSE and clinical diagnosis, data from AIBL (and MACC) was mapped to ADNI space by setting the site *s* to correspond to ADNI.

### 2.8 Dataset prediction model

As one evaluation criterion, we utilized XGBoost to predict which dataset the harmonized brain volumes came from (Figure 2B). The inputs to the XGBoost model were the brain volumes divided by the total intracranial volume (ICV) of each participant. We used logistic regression as the objective function and ensemble of trees as the model structure. Recall that there were 10 groups of harmonized data because of our 10-fold cross-validation procedure (Section 2.4). Therefore, 10 XGBoost classifiers were trained using harmonized MRI volumes from unmatched ADNI and AIBL participants (Figure 2B). For each XGBoost classifier, we used a grid search using the validation group to find the optimal set of hyperparameters.

For evaluation, the 10 XGBoost classifiers were applied to harmonized MRI volumes of matched ADNI and AIBL participants (Figure 2B). The prediction accuracy was averaged across all time points of each participant and the 10 classifiers before averaging across participants. The same procedure was applied to ADNI and MACC participants.

Here, we chose to use XGBoost because it is a powerful classifier for unstructured or tabular data (Grinsztajn et al., 2022; Shwartz-Ziv & Armon, 2022). Using a DNN instead of XGBoost is unlikely to yield very different dataset prediction performance. On the other hand, XGBoost is less sensitive to the choice of hyperparameters compared with DNN, so hyperparameter tuning (and thus training) was a lot faster for XGBoost. Therefore, we chose to use XGBoost for dataset prediction. By contrast, a DNN was utilized for predicting MMSE and clinical diagnosis (i.e., goal-specific DNN), so that the gradients of the goal-specific DNN can be backpropagated to guide the training of the gcVAE model (Section 2.7).

### 2.9 Further analyses

We performed four additional analyses to study the effectiveness of the proposed gcVAE approach.

#### 2.9.1 Effects of training set size

To investigate the effect of training set size on harmonization quality, we repeated the previous analyses (Figures 1 & 2), except that when training harmonization models (Figure 1C), the training set size was varied by sampling 10%, 20%, 30%, 40%, 50%, 60%, 70%, 80 or 90% from the unmatched participants. We repeated this procedure 10 times.

#### 2.9.2 Association analyses

We further investigated the association of the harmonized brain volumes with age, sex, MMSE and clinical diagnosis. We considered all 87 cortical and subcortical gray matter ROIs. For each continuous measure (age or MMSE) and for each ROI, we computed the Pearson’s correlation between the harmonized ROI volume and the continuous measure across matched ADNI and matched AIBL participants. In the case of age, we expected a negative correlation between age and harmonized ROI volumes, so a stronger negative correlation indicates better harmonization. In the case of MMSE, we expect a positive correlation between MMSE and harmonized ROI volumes because lower MMSE indicates greater cognitive decline. Therefore, a greater positive correlation indicates better harmonization. For each discrete variable (clinical diagnosis or sex), we computed *η*^2^ from running ANOVA on the matched ADNI and matched AIBL participants. Greater *η*^2^ indicates greater differences across the groups (e.g., male versus female), suggesting better harmonization. The same procedure was applied to ADNI and MACC.

#### 2.9.3 ComBat with additional covariates

As discussed previously, in our main analyses, we only used age and sex as covariates for the ComBat baseline (Section 2.6.1). Here, we also considered a ComBat variant, where age, sex, MMSE and clinical diagnosis were used as covariates. We note that this version of ComBat assumed that MMSE and clinical diagnosis information were known in the test set (matched participants). Therefore, the prediction performance of ComBat (with the additional covariates) was corrupted by test set leakage and was not valid.

#### 2.9.4 Reversing the roles of the matched and unmatched participants

In the original analyses (Figures 1 and 2), the harmonization models, goal-specific DNNs and dataset prediction models were trained on unmatched participants. The evaluations were then performed on matched participants (Figures 2B and 2C).

In this analysis, we reversed the roles of the matched and unmatched participants (Figures S1 and S2) with two exceptions. First, the prediction performance of unmatched unharmonized ADNI and unmatched unharmonized AIBL participants was not comparable, so the downstream application performance was only evaluated on unmatched unharmonized AIBL and unmatched harmonized AIBL data (compare Figure S2C and Figure 2C).

Second, given the number of matched participants were so small, the training of the goal-specific DNN would not be effective. Therefore, the goal-specific DNN was trained with all (both matched and unmatched) ADNI participants (compare Figure S1B and Figure 1B). We note that this is not an issue since the downstream application performance no longer utilized any ADNI data (Figure S2C).

### 2.10 Deep neural network implementation

All DNNs were implemented using PyTorch (Paszke et al., 2017) and computed on NVIDIA RTX 3090 GPUs with CUDA 11.0. To optimize the DNNs, we used the Adam optimizer (Kingma & Ba, 2017) with default PyTorch settings.

### 2.11 Statistical tests

Two-sided two-sample t-tests were utilized to test for differences in age and MMSE between matched participants of AIBI and ADNI (as well as MACC and ADNI). In the case of sex and clinical diagnoses, we utilized chi-squared tests.

As discussed in Sections 2.5 and 2.8, prediction performance was averaged across all time points of each participant and across the 10 sets of models, yielding a single prediction performance for each participant. Therefore, for each dataset, harmonization approach and evaluation metric, we obtained a performance vector where each element represented one participant. When comparing dataset prediction performance (or goal-specific prediction performance) between two harmonization approaches, a permutation test with 10,000 permutations. Each permutation involves randomly swapping the entries between the performance vectors of the two approaches. Figure S3 illustrates this permutation procedure in more details.

Multiple comparisons were corrected with a false discovery rate (FDR) of q < 0.05.

### 2.12 Data and code availability

Code for the various harmonization algorithms can be found here (GITHUB_LINK). Two co-authors (PC and CZ) reviewed the code before merging it into the GitHub repository to reduce the chance of coding errors.

The ADNI and the AIBL datasets can be accessed via the Image & Data Archive (https://ida.loni.usc.edu/). The MACC dataset can be obtained via a data-transfer agreement with the MACC (http://www.macc.sg/).

## 3 Results

### 3.1 cVAE & gcVAE removed more dataset differences than ComBat

Figure 5A shows the dataset prediction performance for the matched ADNI and AIBL participants. Before harmonization, the XGBoost classifier was able to predict which dataset a participant came from with 100% accuracy. After applying ComBat, the prediction accuracy dropped to 0.626 ± 0.410 (mean ± std), suggesting significant removal of dataset differences. After applying cVAE and gcVAE, dataset prediction performance dropped to 0.595 ± 0.381 and 0.603 ± 0.382 respectively, which were significantly lower than ComBat (Table 3). There was no statistical difference between cVAE and gcVAE. However, dataset prediction accuracies for cVAE and gcVAE were still better than chance (p = 1e-4), suggesting residual dataset differences.

**Figure 5.**
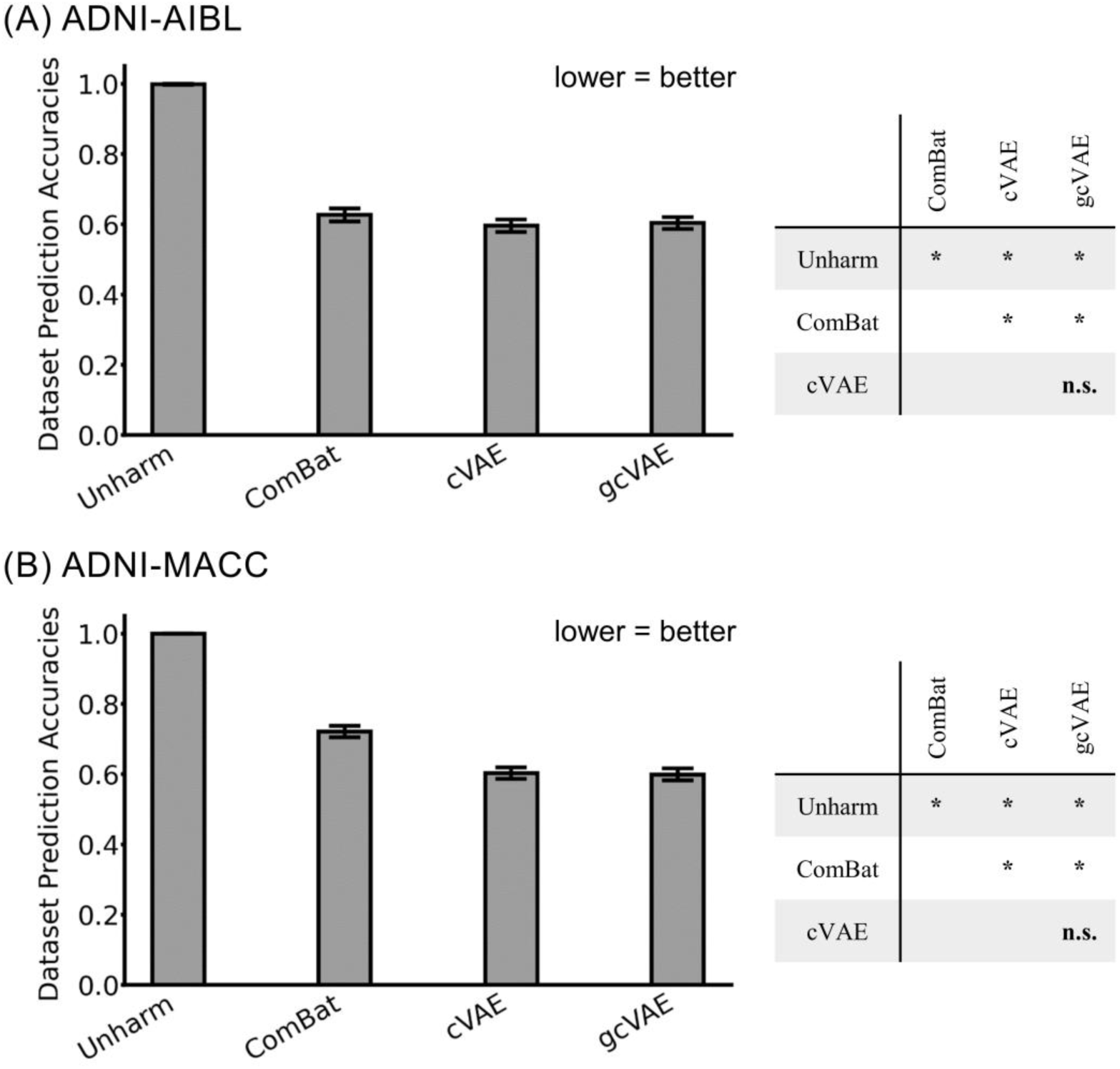
Dataset prediction accuracies. (A) Left: Dataset prediction accuracies for matched ADNI and AIBL participants. Right: p values of differences between different approaches. "*" indicates statistical significance after surviving FDR correction (q < 0.05). "n.s." indicates not significant. (B) Same as (A) but for matched ADNI and MACC participants. All p values are reported in Tables 3 and 4.

**Table 3.**
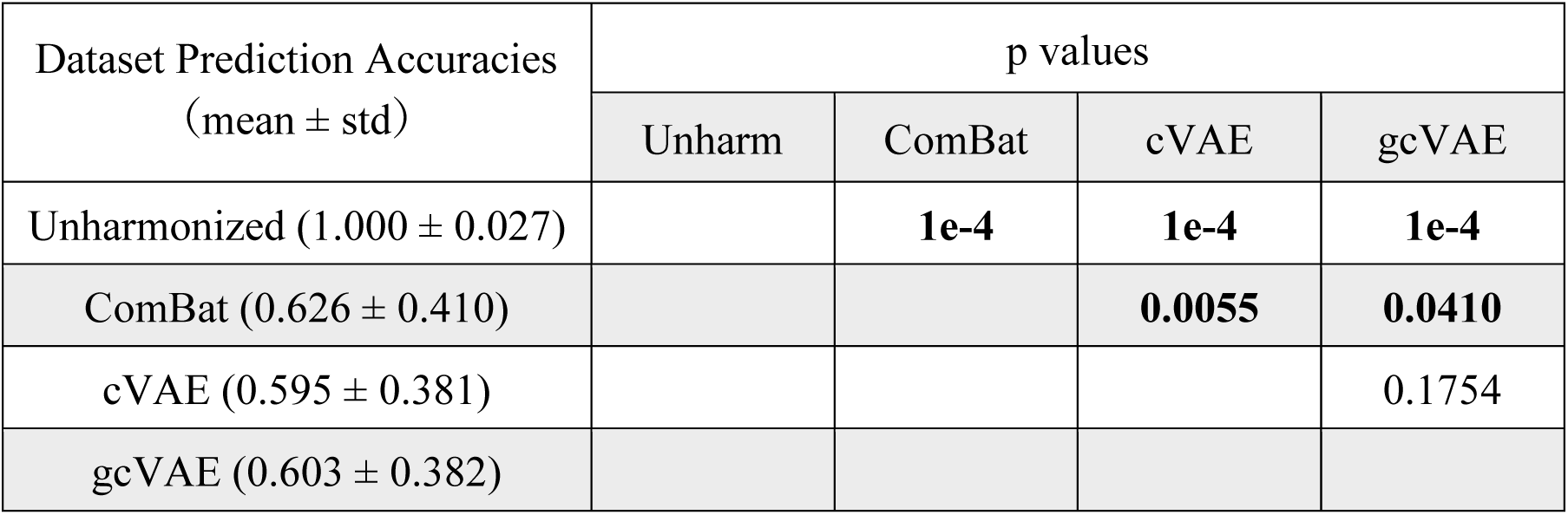
Dataset prediction accuracies with p values of differences between different approaches for matched ADNI and AIBL participants. Statistically significant p values after FDR (q < 0.05) corrections are bolded.

Similar results were obtained for matched ADNI and MACC participants (Figure 5B). Before harmonization, the XGBoost classifier was able to predict which dataset a participant came from with 100% accuracy. Dataset prediction accuracies after ComBat, cVAE and gcVAE were 0.721 ± 0.392, 0.603 ± 0.391 and 0.598 ± 0.398 respectively. There was no statistical difference between cVAE and gcVAE. Both cVAE and gcVAE had statistically lower dataset prediction performance than ComBat (Table 4).

**Table 4.** Dataset prediction accuracies with p values of differences between different approaches for matched ADNI and MACC participants. Statistically significant p values after FDR (q < 0.05) corrections are bolded.

| Dataset Prediction Accuracies<br>(mean ± std) | p values | | | |
| --- | --- | --- | --- | --- |
|  | Unharm | ComBat | cVAE | gcVAE |
| Unharmonized ($1.00 \pm 1e-16$) | | **1e-4** | **1e-4** | **1e-4** |
| ComBat ($0.721 \pm 0.392$) | | | **1e-4** | **1e-4** |
| cVAE ($0.603 \pm 0.391$) | | | | 0.3584 |
| gcVAE ($0.598 \pm 0.398$) | | | | |

Overall, cVAE and gcVAE appeared to remove more dataset differences than ComBat. However, dataset prediction accuracies for cVAE and gcVAE were still better than chance (p = 1e-4), suggesting residual dataset differences.

### 3.2 gcVAE outperformed cVAE for clinical diagnosis prediction

Figure 6A shows the clinical diagnosis prediction accuracies for matched ADNI and AIBL participants. Because the matched participants had similar age, sex, MMSE and clinical diagnosis (Figure 3), comparison between unharmonized ADNI and unharmonized AIBL participants would indicate whether there was a drop in prediction performance due to dataset differences (e.g., scanner differences). Unexpectedly, there was no statistical difference in clinical diagnosis prediction performance between unharmonized ADNI and unharmonized AIBL participants (Table 5).

**Figure 6.**
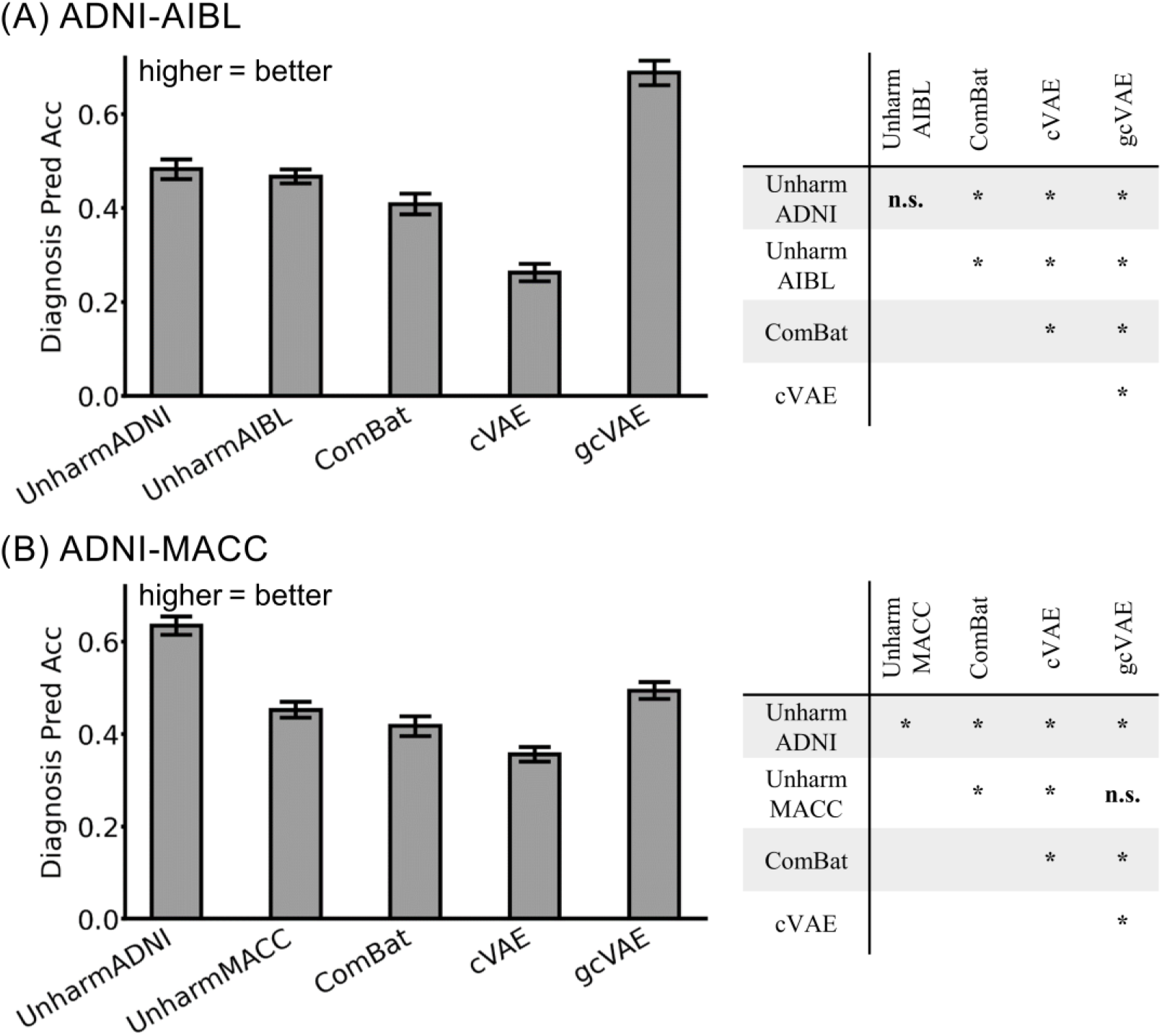
Clinical diagnosis prediction accuracies. (A) Left: Clinical diagnosis prediction accuracies for matched ADNI and AIBL participants. Right: p values of differences between different approaches. "*" indicates statistical significance after surviving FDR correction (q < 0.05). "n.s." indicates not significant. (B) Same as (A) but for matched ADNI and MACC participants. All p values are reported in Tables 5 and 6.

**Table 5.** Clinical diagnosis prediction accuracies with p values of differences between different approaches for matched ADNI and AIBL participants. Statistically significant p values after FDR (q < 0.05) corrections are bolded.

| Clinical Diagnosis Prediction<br>Accuracies (mean ± std) | p values | | | | |
| --- | --- | --- | --- | --- | --- |
|  | Unharm ADNI | Unharm AIBL | ComBat | cVAE | gcVAE |
| Unharm ADNI (0.48 ± 0.33) | | 0.5171 | **0.0077** | **1e-4** | **2e-4** |
| Unharm AIBL (0.47 ± 0.23) | | | **7e-4** | **1e-4** | **1e-4** |
| ComBat (0.41 ± 0.34) | | | | **1e-4** | **1e-4** |
| cVAE (0.26 ± 0.29) | | | | | **1e-4** |
| gcVAE (0.69 ± 0.41) | | | | | |

Applying ComBat resulted in a statistically significant drop in prediction performance (p = 7e-4) compared with no harmonization. This suggests that ComBat removed biological information in addition to dataset differences (Figure 5A). cVAE exhibited an even bigger drop in prediction performance compared with ComBat (p = 1e-4), suggesting that the better removal of dataset differences (Figure 5A) came at the expense of removing even more biological information. gcVAE yielded the best prediction performance with statistically significant improvements over all other approaches, including unharmonized ADNI (see p values in Table 5).

Figure 6B shows the clinical diagnosis prediction accuracies for matched ADNI and MACC participants. As expected, there was a significant drop in clinical diagnosis prediction performance between unharmonized ADNI and unharmonized MACC participants (p = 1e-4). The decrease in clinical diagnosis performance was worsened by ComBat and cVAE, once again suggesting that the removal of dataset differences (Figure 5B) came at the expense of also removing biological information. gcVAE recovered a significant portion of the decrease in prediction performance, such that it was not statistically different from unharmonized MACC (Table 6). However, it was still significantly worse than unharmonized ADNI, suggesting potential room for improvement.

**Table 6.** Clinical diagnosis prediction accuracies with p values of differences between different approaches for matched ADNI and MACC participants. Statistically significant p values after FDR (q < 0.05) corrections are bolded.

| Clinical Diagnosis Prediction<br>Accuracies (mean ± std) | p values | | | | |
| --- | --- | --- | --- | --- | --- |
|  | Unharm ADNI | Unharm MACC | ComBat | cVAE | gcVAE |
| Unharm ADNI (0.63 ± 0.33) | | **1e-4** | **1e-4** | **1e-4** | **1e-4** |
| Unharm MACC (0.45 ± 0.29) | | | **0.0124** | **1e-4** | 0.0545 |
| ComBat (0.42 ± 0.35) | | | | **2e-4** | **0.0065** |
| cVAE (0.36 ± 0.26) | | | | | **1e-4** |
| gcVAE (0.49 ± 0.30) | | | | | |

### 3.3 gcVAE outperformed cVAE in MMSE prediction

Figure 7A shows the MMSE prediction mean absolute error (MAE) for matched ADNI and AIBL participants. Because the matched participants had similar age, sex, MMSE and clinical diagnosis, comparison between unharmonized ADNI and unharmonized AIBL participants would indicate whether there was a drop in prediction performance due to dataset differences (e.g., scanner differences). As expected, there was a drop in MMSE prediction performance (increased MAE) for unharmonized AIBL participants compared with unharmonized ADNI participants (p = 1e-4).

**Figure 7.**
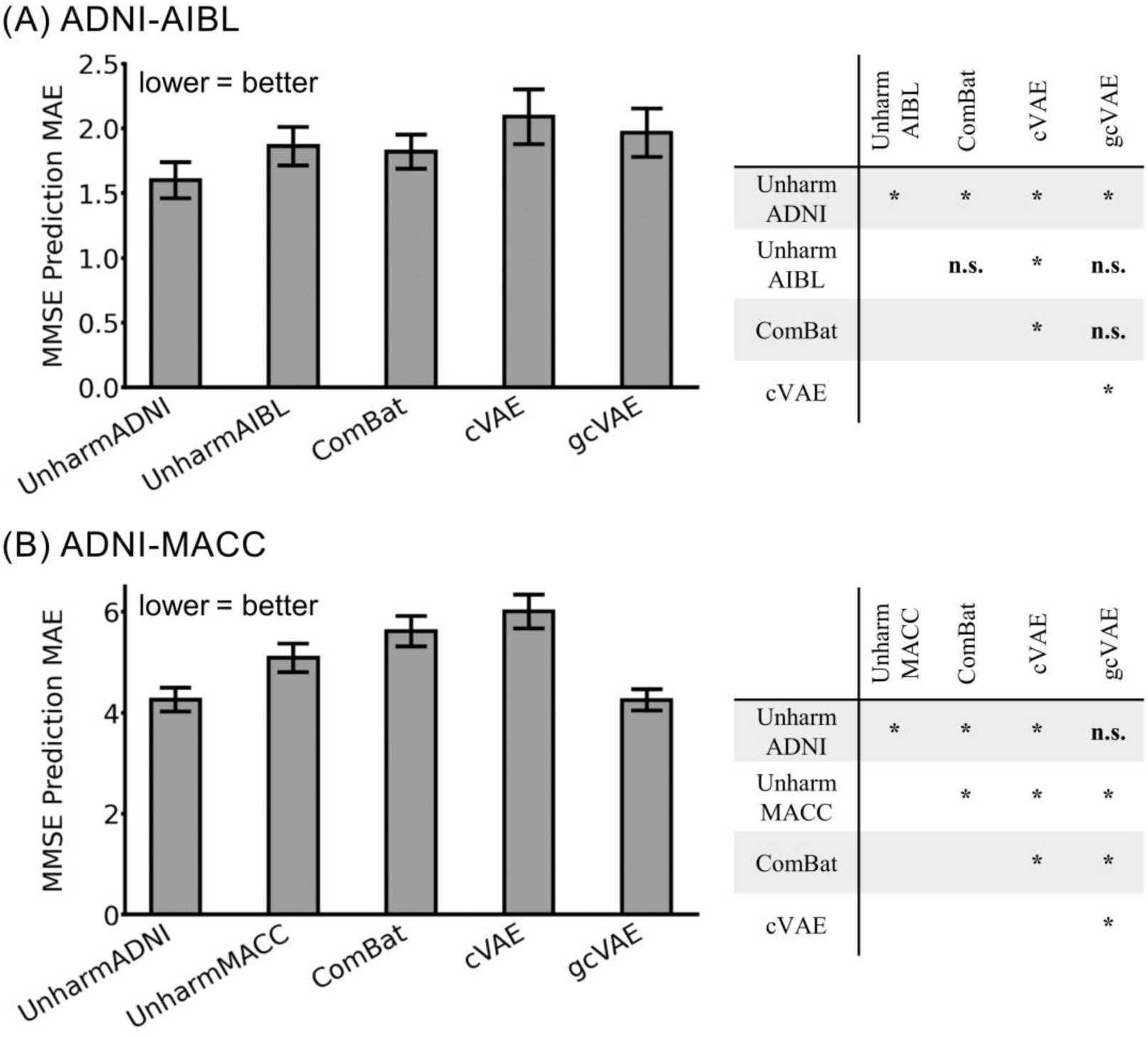
MMSE prediction errors as measured by mean absolute error (MAE). (A) Left: MMSE prediction errors for matched ADNI and AIBL participants. Right: p values of differences between different approaches. "*" indicates statistical significance after surviving FDR correction (q < 0.05). "n.s." indicates not significant. (B) Same as (A) but for matched ADNI and MACC participants. All p values are reported in Tables 7 and 8.

**Table 7.** MMSE prediction errors with p values of differences between different approaches for matched ADNI and AIBL participants. Statistically significant p values after FDR (q < 0.05) corrections are bolded.

| MMSE Prediction MAE<br>(mean ± std) | p values | | | | |
| --- | --- | --- | --- | --- | --- |
|  | Unharm ADNI | Unharm AIBL | ComBat | cVAE | gcVAE |
| Unharm ADNI (1.60 ± 2.17) | | **1e-4** | **0.0061** | **1e-4** | **1e-4** |
| Unharm AIBL (1.86 ± 2.32) | | | **0.4339** | **0.0054** | 0.0756 |
| ComBat (1.82 ± 2.07) | | | | **0.0322** | 0.1473 |
| cVAE (2.09 ± 3.29) | | | | | **0.0023** |
| gcVAE (1.97 ± 2.93) | | | | | |

There was no statistical difference between ComBat and the unharmonized AIBL participants. cVAE had statistically worse MMSE prediction performance compared with all other approaches (p values in Table 7). gcVAE recovered a significant portion of the decrease in prediction performance, such that prediction performance was not statistically different from ComBat and unharmonized AIBL (Table 7). However, it was still statistically worse than unharmonized ADNI, suggesting further room for improvement.

Figure 7B shows the MMSE prediction MAE for matched ADNI and MACC participants. As expected, there was a drop in MMSE prediction performance (increased MAE) for unharmonized MACC participants compared with unharmonized ADNI participants (p = 1e-4). Both ComBat and cVAE caused further drop in prediction performance (p values in Table 8). gcVAE had the best prediction performance (lowest MAE), such that prediction performance was statistically better than unharmonized MACC and not statistically different from unharmonized ADNI (Table 8).

**Table 8.** MMSE prediction errors with p values of differences between different approaches for matched ADNI and MACC participants. Statistically significant p values after FDR (q < 0.05) corrections are bolded.

| MMSE Pred MAE<br>(mean ± std) | p values | | | | |
| --- | --- | --- | --- | --- | --- |
|  | Unharm ADNI | Unharm MACC | ComBat | cVAE | gcVAE |
| Unharm ADNI (4.26 ± 3.87) | | **1e-4** | **1e-4** | **1e-4** | 0.9570 |
| Unharm MACC (5.09 ± 4.66) | | | **1e-4** | **1e-4** | **1e-4** |
| ComBat (5.61 ± 5.03) | | | | **1e-4** | **1e-4** |
| cVAE (5.96 ± 5.50) | | | | | **1e-4** |
| gcVAE (4.25 ± 3.57) | | | | | |

### 3.4 Further analyses

#### 3.4.1 Effects of training set size

We investigated the effects of varying the training set size used for fitting the harmonization models (Figure 1C). Across all sample sizes, cVAE and gcVAE generally achieved lower dataset prediction accuracies than ComBat (Figure 8A). Across all sample sizes, gcVAE achieved better clinical diagnosis prediction than cVAE and ComBat (Figure 8B). Across all sample sizes for MACC, gcVAE achieved better MMSE prediction than cVAE and ComBat (Figure 8C2). Across all sample sizes for AIBL (Figure 8C1), gcVAE achieved better MMSE prediction than cVAE; gcVAE achieved worse prediction than ComBat. Overall, across all sample sizes, gcVAE compared favorably with cVAE and ComBat.

**Figure 8.**
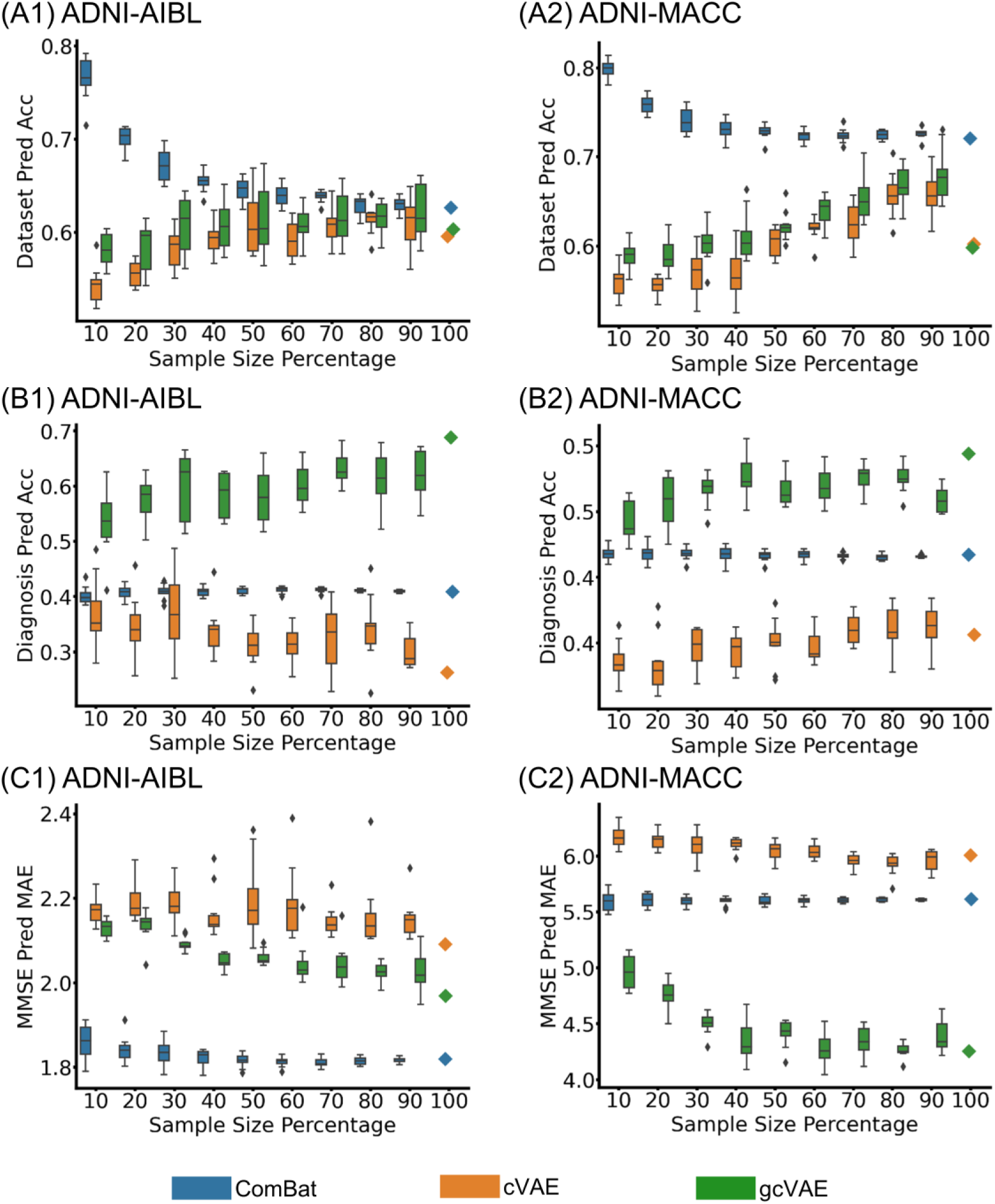
Performance of harmonization models trained with different sample sizes. (A1) Dataset prediction accuracies for matched ADNI-AIBL participants; (B1) The clinical diagnosis prediction accuracies for matched AIBL participants; (C1) MMSE prediction errors for matched AIBL participants. (A2), (B2), and (C2) are the same as (A1), (B1), and (C1), but for matched ADNI and MACC participants.

In the case of ComBat, larger sample sizes led to worse dataset prediction accuracies (i.e., better harmonization). However, sample sizes have minimal effect on clinical diagnosis and MMSE prediction. In the case of cVAE, greater sample sizes led to better MMSE prediction for both AIBL and MACC participants, better clinical diagnosis prediction for MACC participants, worse clinical diagnosis prediction for AIBL participants, and better dataset prediction accuracies. In the case of gcVAE, greater sample sizes led to better MMSE and clinical diagnosis prediction for both AIBL and MACC participants, as well as better dataset prediction accuracies. Overall, for both cVAE and gcVAE, larger sample sizes appeared to improve downstream application performance (i.e., MMSE and clinical diagnosis prediction), but at the expense of dataset prediction performance.

#### 3.4.2 Association analyses

Figures 9 shows the association analyses between gray matter ROI volumes and four variables (age, sex, MMSE and clinical diagnoses) among matched ADNI and AIBL participants. Figure 10 shows the same analyses for matched ADNI and MACC participants. For each scatter plot, more dots in the green region indicates better gcVAE performance compared with the baseline. gcVAE clearly outperformed no harmonization (Figures 9A and 10A) and ComBat (Figures 9B and 10B) in both datasets. On the other hand, cVAE and gcVAE exhibited comparable performance (Figures 9C and 10C) in both datasets.

**Figure 9.**
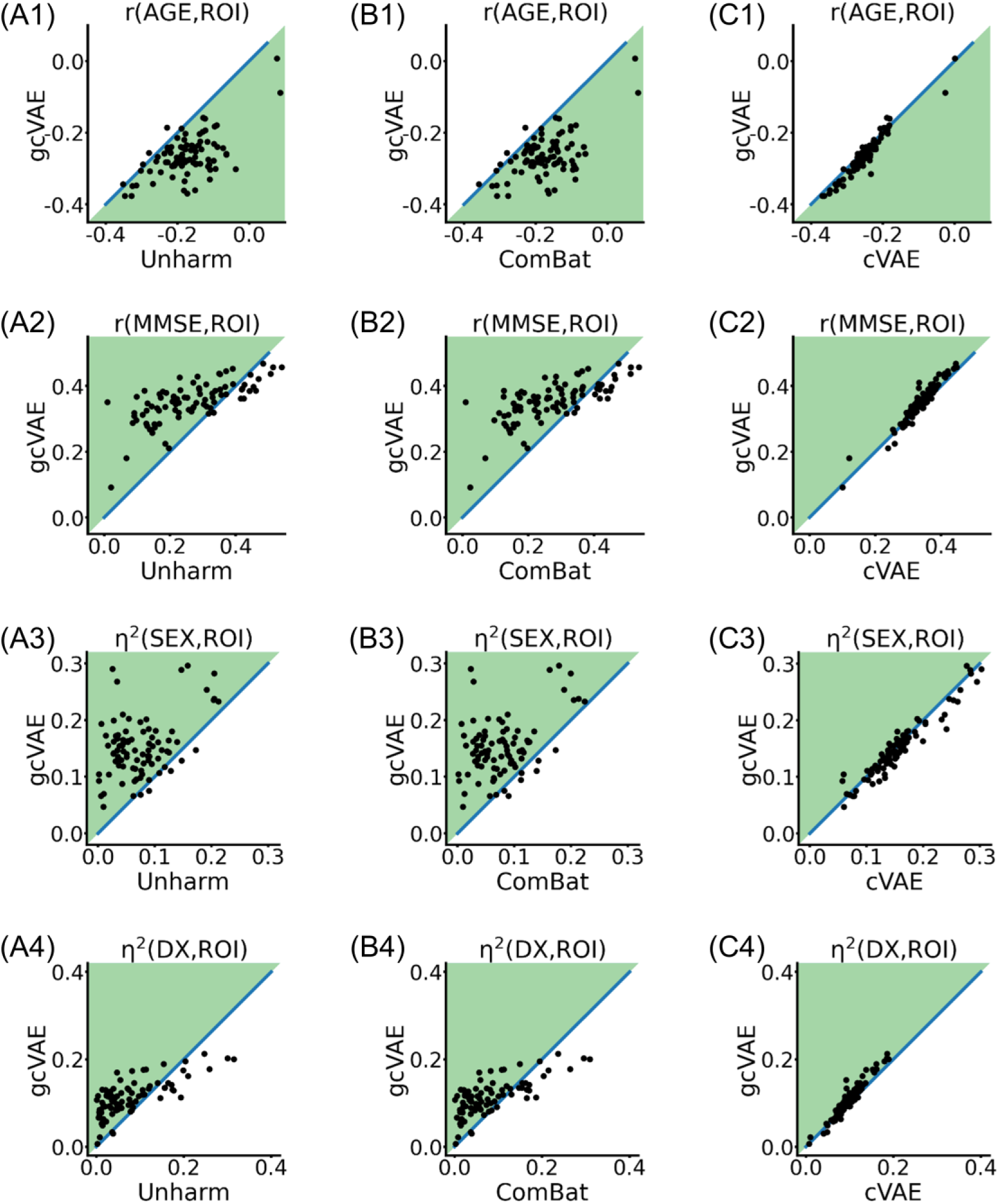
Association analyses between gray matter ROI volumes and four variables (age, MMSE, sex and clinical diagnosis) for matched ADNI and AIBL participants. First row shows association with age. Second row shows association with MMSE. Third row shows association with sex. Fourth row shows association with clinical diagnosis. (A) Comparison between gcVAE and no harmonization. (B) Comparison between gcVAE and ComBat. (C) Comparison between gcVAE and cVAE. Each block dot represents one gray matter ROI. Dots in the green area indicates better gcVAE performance compared with baseline. gcVAE clearly outperforms no harmonization and ComBat. gcVAE and cVAE exhibited similar performance.

**Figure 10.**
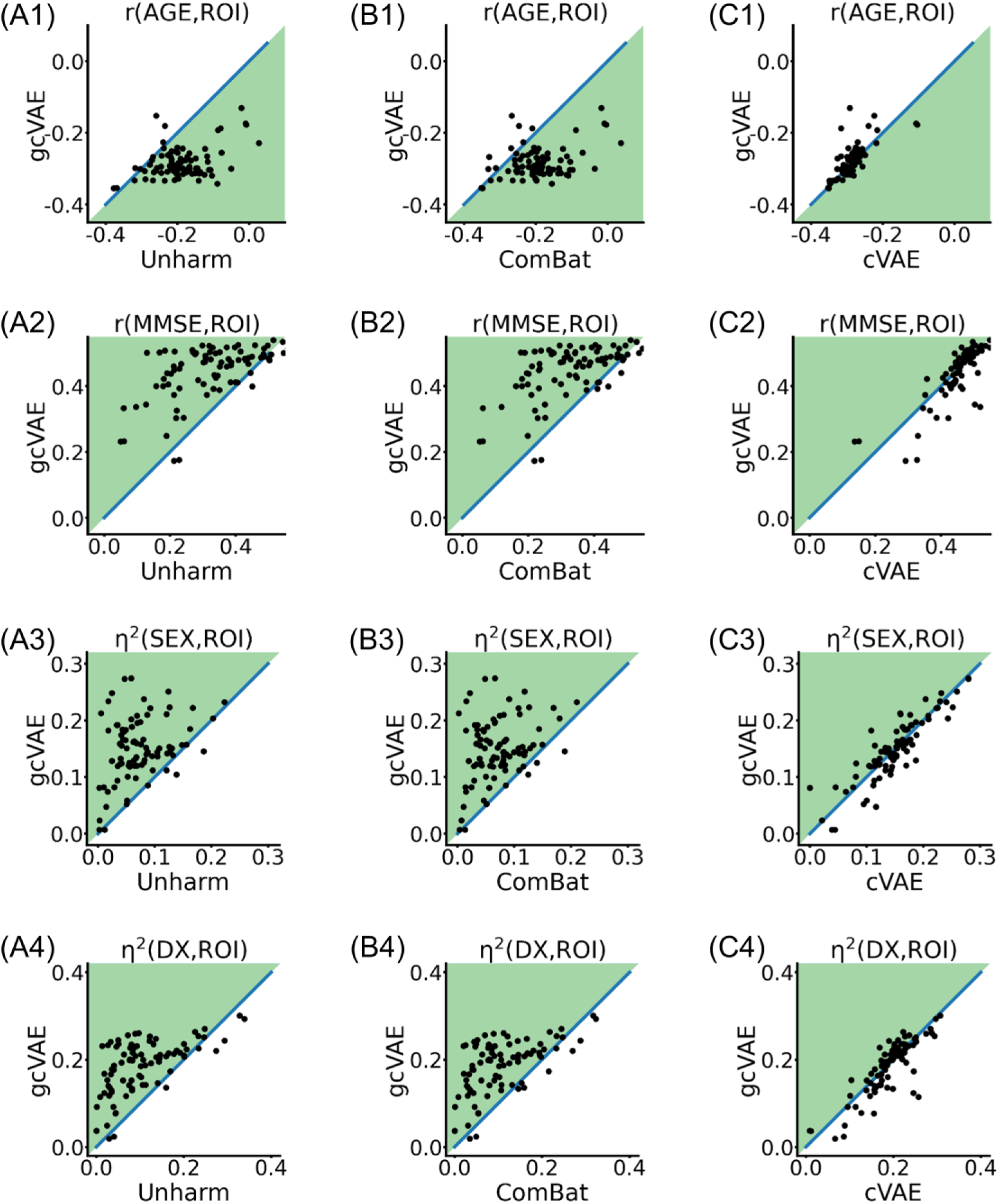
Association analyses between gray matter ROI volumes and four variables (age, MMSE, sex and clinical diagnosis) for matched ADNI and MACC participants. First row shows association with age. Second row shows association with MMSE. Third row shows association with sex. Fourth row shows association with clinical diagnosis. (A) Comparison between gcVAE and no harmonization. (B) Comparison between gcVAE and ComBat. (C) Comparison between gcVAE and cVAE. Each block dot represents one gray matter ROI. Dots in the green area indicates better gcVAE performance compared with baseline. gcVAE clearly outperforms no harmonization and ComBat. gcVAE and cVAE exhibited similar performance.

#### 3.4.3 ComBat with additional covariates

Our main analysis utilized ComBat with age and sex as covariates. Here, we considered ComBat with age, sex, MMSE and clinical diagnosis as covariates. We note that this version of ComBat assumed that MMSE and clinical diagnosis information were known in the test set (matched participants). Therefore, the prediction performance of ComBat (with the additional covariates) was corrupted by test set leakage and was not valid.

The additional covariates led to better MMSE and clinical diagnosis prediction by ComBat (Tables S10 and S11). In the case of AIBL, clinical diagnosis prediction remained statistically worse than gcVAE, but MMSE prediction was now statistically better than gcVAE. In the case of MACC, clinical diagnosis prediction was now comparable with gcVAE, but MMSE prediction remained worse than gcVAE. Interestingly, the additional covariates led to greater dataset prediction accuracies for both ADNI-AIBL and ADNI-MACC, suggesting worse harmonization. Together, gcVAE remained better than ComBat.

#### 3.4.4 Reversing the roles of the matched and unmatched participants

In this analysis, we reversed the roles of matched and unmatched participants (Section 2.9.4). Similar to the original main analyses, we found that gcVAE compared favorably with both ComBat and cVAE (Figures S4 to S6; Tables S12 to S17).

More specifically, recall that there were six evaluation metrics (two for dataset prediction, two for diagnosis prediction and two for MMSE prediction). gcVAE was statistically better than ComBat for both dataset prediction metrics and two downstream application performance metrics, while being statistically worse than ComBat in one downstream application performance metric (Figures S4 to S6; Tables S12 to S17). On the other hand, gcVAE was statistically worse than cVAE for the two dataset prediction metrics, but statistically better than cVAE for the four downstream application performance metrics (Figures S4 to S6; Tables S12 to S17). Therefore, similar to the main results, cVAE removed more dataset differences at the expense of removing more biological information.

One interesting deviation from the main results was that in the current setup (where harmonization models were trained on matched participants), ComBat was statistically better than no harmonization across all six evaluation metrics. On the other hand, in the main analysis (Figures 5 to 7; Tables 3 to 8), ComBat was statistically better than no harmonization for both dataset prediction metrics, but statistically worse than no harmonization for all four downstream application performance metrics. On the other hand, for the main analysis, gcVAE was statistically better than no harmonization for both dataset prediction metrics and two downstream application performance metrics. In the current analysis, gcVAE was statistically better than no harmonization for both dataset prediction metrics and three downstream application performance metrics, but was statistically worse for one application performance metric. Therefore, gcVAE appeared more robust than ComBat to covariate differences during the harmonization procedure.

## 4 Discussion

In this study, we proposed a flexible harmonization framework to utilize downstream application performance to regularize the harmonization model. Our proposed approach could be integrated with most harmonization approaches based on DNNs. Here, we integrated our approach with the cVAE model. Using three large-scale datasets, we demonstrated that gcVAE compared favorably with ComBat and cVAE.

We found that cVAE was able to significantly remove more dataset differences than ComBat (Figure 5). This makes intuitive sense given that cVAE considered all brain regions jointly, so should theoretically be able to remove multivariate site effects distributed across brain regions. However, the removal of more dataset differences came at the expense of also removing relevant biological information as measured by downstream application performance (Figures 6 and 7).

Indeed, the removal of relevant biological information was an issue not just for cVAE, but also for ComBat. In the case of predicting clinical diagnosis and MMSE, the use of ComBat led to similar or worse performance than not harmonizing at all. By constraining the harmonization with goal-specific DNNs, the gcVAE models were able to yield better prediction of MMSE and clinical diagnosis (Figures 6 and 7), while removing as much dataset differences as cVAE (Figure 5).

In the case of clinical diagnosis prediction, gcVAE was able to yield better prediction performance than no harmonization. In the case of MMSE prediction, gcVAE was able to yield better prediction performance than no harmonization in the MACC dataset, but was only able to yield comparable prediction performance than no harmonization in the AIBL dataset.

Our main analyses (Figures 6 and 7) showed that gcVAE facilitated the translation of goal-specific DNNs from ADNI to new datasets (AIBL and MACC). Another common application of harmonization is to facilitate the pooling of datasets for some joint analysis.

Here, we investigated the association of the brain volumes with multiple variables across the harmonized datasets. We found that gcVAE clearly outperformed no harmonization (Figures 9A and 10A) and ComBat (Figures 9B and 10B). On the other hand, cVAE and gcVAE exhibited comparable performance (Figures 9C and 10C).

### 4.1 Matched versus unmatched participants

We note that our workflow utilized unmatched participants to train the harmonization models, dataset prediction models and goal-specific DNNs, while evaluation was performed on the matched participants (Figures 1 and 2). The setup allowed us to compare downstream application performance between unharmonized data from matched ADNI participants and matched AIBL participants. Because age, sex, MMSE and clinical diagnosis were similar between matched ADNI and AIBL participants, the drop in downstream application performance (clinical diagnosis or MMSE prediction) could be attributed to a lack of harmonization. Since the goal-specific DNNs were trained on ADNI (Figure 2B), the prediction performance on matched unharmonized ADNI participants served as an upper bound on the prediction performance after harmonization.

Surprisingly, in the case of clinical diagnosis prediction in the AIBL dataset, gcVAE was better than the upper bound (Figure 6A). On the other hand, in the case of MMSE prediction in the AIBL dataset, gcVAE only achieved similar performance as no harmonization and was worse than the upper bound (Figure 7A). One possible reason for this discrepancy is that when tuning the hyperparameters, the weights tradeoff the prediction of MMSE and clinical diagnosis were fixed, so in the case of AIBL, this might have inadvertently favored clinical diagnosis prediction more than MMSE prediction.

However, we note that the current workflow of training on unmatched participants can prove challenging for ComBat (Nygaard et al., 2016; Zindler et al., 2020) because of covariate differences between ADNI and AIBL (as well ADNI and MACC). Therefore, we considered a control analysis in which the roles of the matched and unmatched participants were swapped. Consistent with the main analyses, we found that gcVAE compared favorably with both ComBat and cVAE (Figures S4 to S6; Tables S12 to S17). Furthermore, in the control analysis, ComBat was better than no harmonization for both dataset prediction and downstream application performance. On the other hand, in the control analysis, gcVAE was statistically better than no harmonization for both dataset prediction metrics and three downstream application performance metrics, but was statistically worse for one application performance metric. Overall, this suggests that gcVAE was more robust than ComBat to covariate differences between datasets used for the harmonization procedure.

### 4.2 Sample size

Deep neural networks are often thought to be data hungry. Across different sample sizes (Figure 8), gcVAE was better than cVAE for all four downstream application performance. On the other hand, across all sample sizes, gcVAE was better than ComBat for three downstream prediction metrics. Interestingly, gcVAE was worse than ComBat for MMSE prediction in the AIBL dataset across all sample sizes but given the rapid improvement trajectory of gcVAE (Figure 8C1), we might expect the gap to close rapidly with more data. Surprisingly, as the sample size increases, the downstream performance of gcVAE improved at the expense of dataset prediction performance. However, the dataset prediction accuracies of gcVAE continued to be worse (i.e., better harmonization) than ComBat even with the full set of data (Figure 8A).

### 4.3 Methodological considerations

To illustrate the use of gcVAE, when harmonizing ADNI and a new dataset, the researcher could validate gcVAE by repeating the same procedure as the current study (Figures 2 and 3). Once the researcher is satisfied with the performance, the researcher could then train the model on 90% of the data and tune the hyperparameters on the remaining 10% of the data without the need of a 10-fold cross-validation procedure. The final model can then be applied to the full dataset.

An interesting methodological consideration is the handling of confound variables when using gcVAE. For example, age is likely related to clinical diagnosis. Therefore, when training gcVAE to harmonize ADNI and AIBL, the algorithm might seek to preserve age-related brain patterns related to clinical diagnosis. However, we note that this may or may not be an issue depending on the study. For example, if our goal is clinical diagnosis, then it would be counterproductive to exclude age in the diagnosis procedure. After all, demographics are often used for differential diagnosis in actual clinical practice.

There might indeed be situations, where the related variables are indeed confounds. For example, if a study is interested in dementia risks above and beyond aging, then age does become a confound. In that scenario, researchers could consider regressing age from the imaging features and/or target variables before training the goal-specific DNN. Another approach is to include an adversarial cost when training the goal-specific DNN to ensure the intermediate layers could not be used to predict the confound variable (e.g., age).

The theoretical advantage of gcVAE over ComBat is its multivariate nature, which allowed cVAE to remove site differences distributed across brain regions. This advantage is clearly demonstrated in the dataset prediction experiments (Figure 5). More recent ComBat variants, such as CovBat (Chen et al., 2019) allowed the harmonization of inter-regional covariance. Given their multivariate nature, cVAE and gcVAE should also in principle remove site variation in covariance.

Finally, our current study only demonstrated results from harmonizing pairs of datasets (ADNI and AIBL, as well as ADNI and MACC). However, the cVAE framework is highly flexible and the cVAE machinery can be easily extended to multiple datasets. Similarly, the goal-specific DNN could also be trained on multiple datasets. So overall, gcVAE could in principle be applied to harmonize multiple datasets jointly. However, this is not something we have demonstrated in this study, which we leave for future work.

### 4.4 Limitations

The strength of gcVAE is also its main limitation. The reliance of goal-specific DNNs led to better downstream performance, but the resulting improvements might not generalize to new downstream applications. Therefore, the training procedure might have to be repeated for each new downstream application. Future research is necessary to address this limitation.

## 5 Conclusion

In this study, we proposed a goal-specific brain MRI harmonization framework, which took into account downstream application performance in the harmonization process. Using three large-scale datasets, we demonstrated that our approach compared favorably with existing approaches in terms of preserving relevant biological information, while removing site differences.

## Acknowledgment

Our research is currently supported by the Singapore National Research Foundation (NRF) Fellowship (Class of 2017), the NUS Yong Loo Lin School of Medicine (NUHSRO/2020/124/TMR/LOA), the Singapore National Medical Research Council (NMRC) LCG (OFLCG19May-0035), NMRC STaR (STaR20nov-0003), and the USA NIH (R01MH120080). Our computational work was partially performed on resources of the National Supercomputing Centre, Singapore (https://www.nscc.sg). Any opinions, findings and conclusions or recommendations expressed in this material are those of the authors and do not reflect the views of the Singapore NRF or the Singapore NMRC. Data collection and sharing for this project was funded by the Alzheimer’s Disease Neuroimaging Initiative (ADNI) (National Institutes of Health Grant U01 AG024904) and DOD ADNI (Department of Defense award number W81XWH-12-2-0012). ADNI is funded by the National Institute on Aging, the National Institute of Biomedical Imaging and Bioengineering, and through generous contributions from the following: AbbVie, Alzheimer’s Association; Alzheimer’s Drug Discovery Foundation; Araclon Biotech; BioClinica, Inc.; Biogen; Bristol-Myers Squibb Company; CereSpir, Inc.; Cogstate; Eisai Inc.; Elan Pharmaceuticals, Inc.; Eli Lilly and Company; EuroImmun; F. Hoffmann-La Roche Ltd and its affiliated company Genentech, Inc.; Fujirebio; GE Healthcare; IXICO Ltd.;Janssen Alzheimer Immunotherapy Research & Development, LLC.; Johnson & Johnson Pharmaceutical Research & Development LLC.; Lumosity; Lundbeck; Merck & Co., Inc.;Meso Scale Diagnostics, LLC.; NeuroRx Research; Neurotrack Technologies; Novartis Pharmaceuticals Corporation; Pfizer Inc.; Piramal Imaging; Servier; Takeda Pharmaceutical Company; and Transition Therapeutics. The Canadian Institutes of Health Research is providing funds to support ADNI clinical sites in Canada. Private sector contributions are facilitated by the Foundation for the National Institutes of Health (www.fnih.org). The grantee organization is the Northern California Institute for Research and Education, and the study is coordinated by the Alzheimer’s Therapeutic Research Institute at the University of Southern California. ADNI data are disseminated by the Laboratory for Neuro Imaging at the University of Southern California.

## Supplementary Material

**Table S1.**
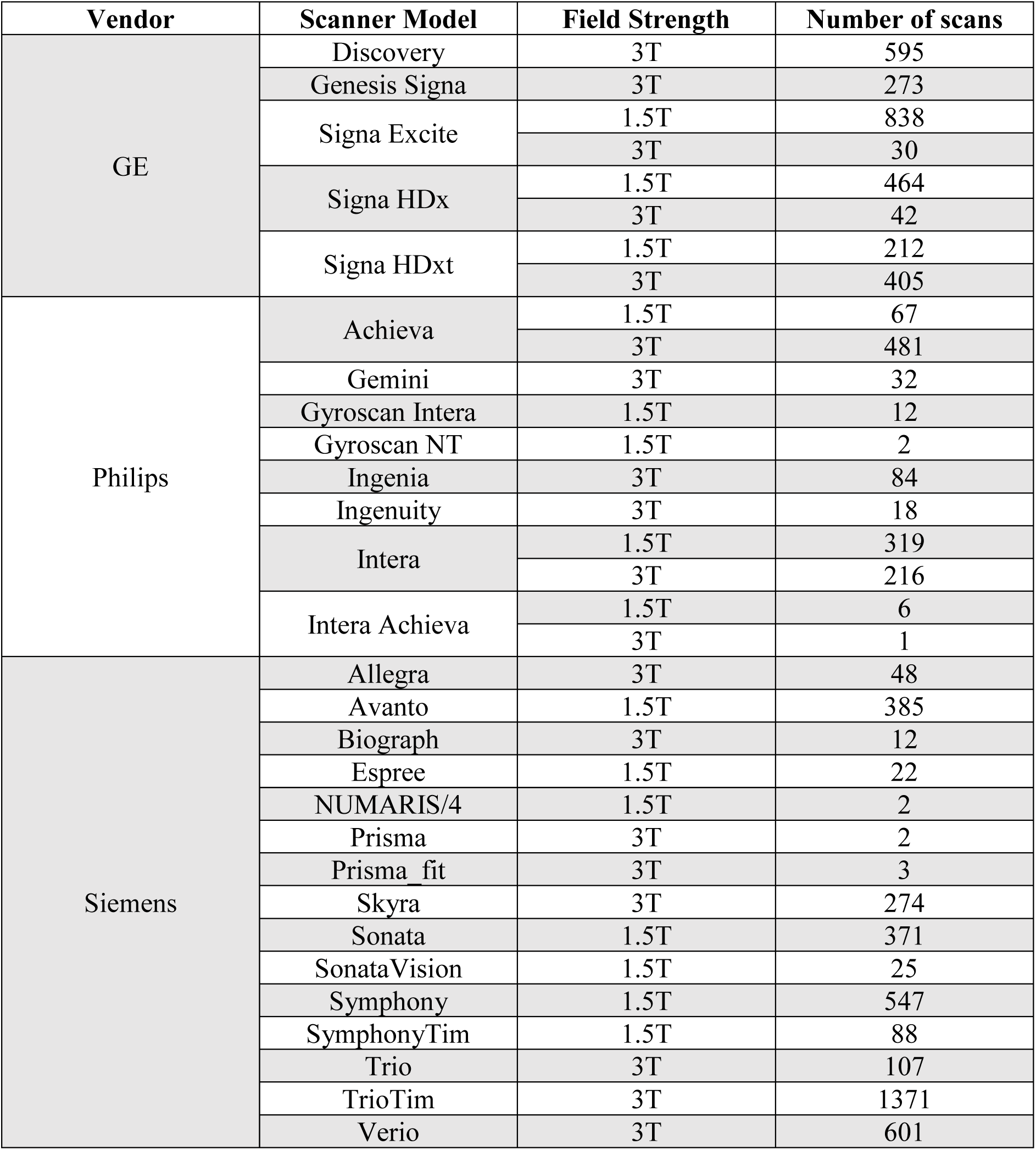
**Scanner information for 7955 scans in ADNI dataset.**

**Table S2.** **Scanner information for 933 scans in AIBL dataset.**

| Vendor | Scanner Model | Field Strength | Number of scans |
| --- | --- | --- | --- |
| Siemens | Avanto | 1.5T | 241 |
|  | TrioTim | 3T | 558 |
|  | Verio | 3T | 134 |

**Table S3.**
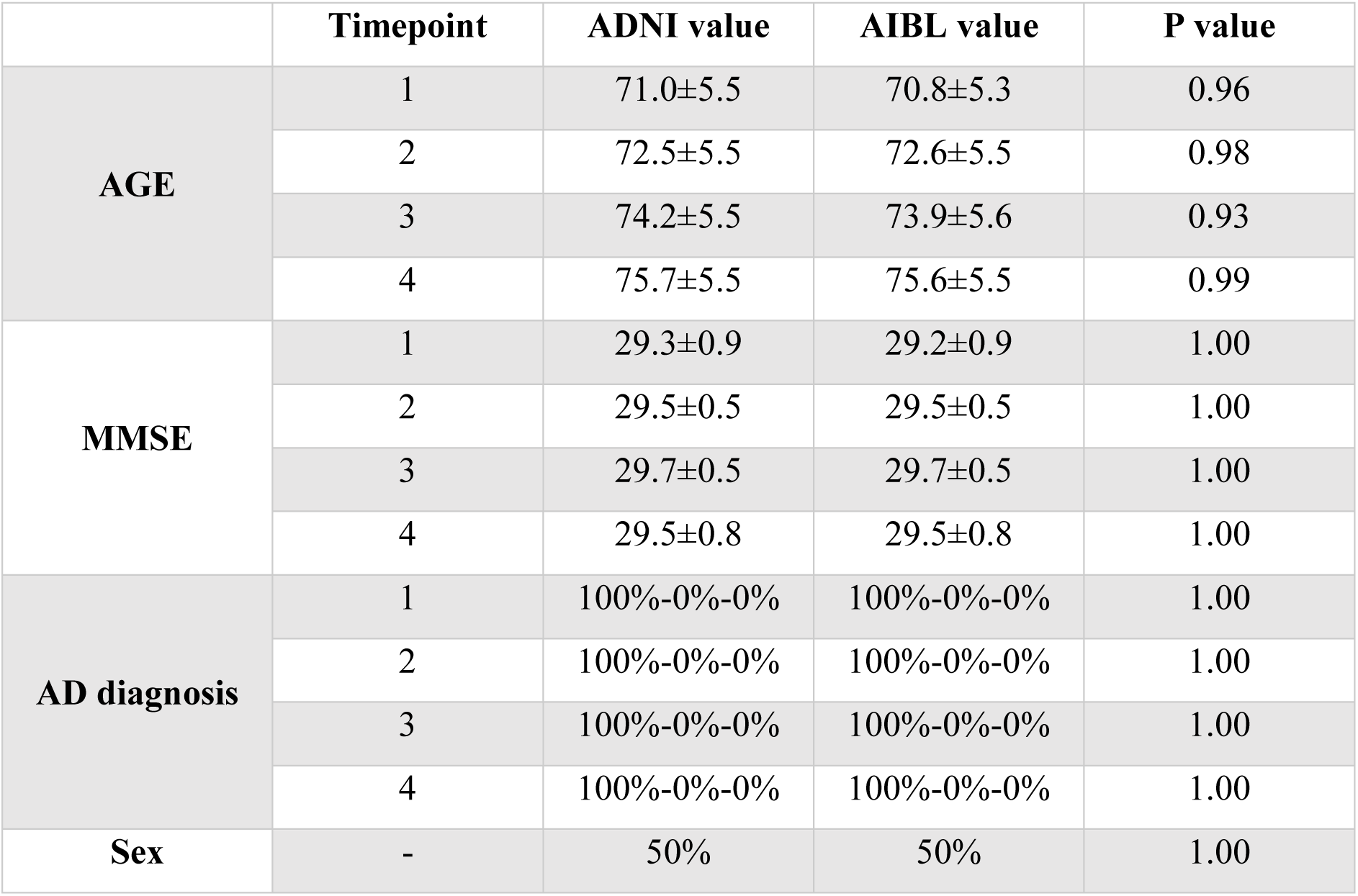
ADNI-AIBL matching results for participants having 4 time points (scans). For clinical diagnosis in the table, the percentage is showed as CN%-MCI%-AD%. For sex in the table, the portion is the ratio of male subjects. For Age/MMSE, the p value was calculated from a two-sample t-test. For Sex/AD diagnosis, the p value was calculated from the chi-square goodness of fit test.

**Table S4.** ADNI-AIBL matching results for participants having 3 time points (scans). For clinical diagnosis in the table, the percentage is showed as CN%-MCI%-AD%. For sex in the table, the portion is the ratio of male subjects. For Age/MMSE, the p value was calculated from a two-sample t-test. For Sex/AD diagnosis, the p value was calculated from the chi-square goodness of fit test.

|  | Timepoint | ADNI value | AIBL value | P value |
| --- | --- | --- | --- | --- |
| AGE | 1 | 73.3±3.3 | 73.1±3.3 | 0.96 |
|  | 2 | 74.8±3.3 | 75.2±3.3 | 0.94 |
|  | 3 | 76.3±3.3 | 76.1±3.3 | 0.97 |
| MMSE | 1 | 29.0±0.0 | 20.0±0.0 | 1.00 |
|  | 2 | 30.0±0.0 | 30.0±0.0 | 1.00 |
|  | 3 | 30.0±0.0 | 30.0±0.0 | 1.00 |
| AD diagnosis | 1 | 100%-0%-0% | 100%-0%-0% | 1.00 |
|  | 2 | 100%-0%-0% | 100%-0%-0% | 1.00 |
|  | 3 | 100%-0%-0% | 100%-0%-0% | 1.00 |
| Sex | - | 50% | 50% | 1.00 |

**Table S5.**
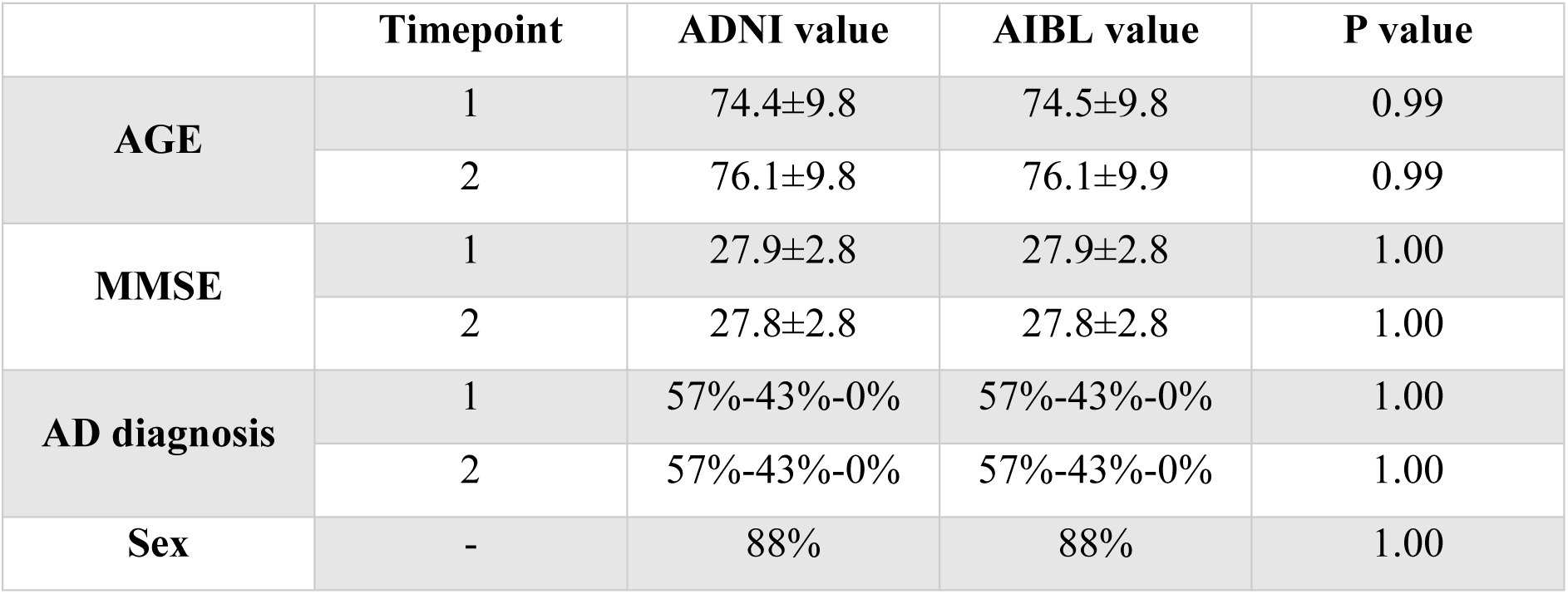
ADNI-AIBL matching results for participants having 2 time points (scans). For clinical diagnosis in the table, the percentage is showed as CN%-MCI%-AD%. For sex in the table, the portion is the ratio of male subjects. For Age/MMSE, the p value was calculated from a two-sample t-test. For Sex/AD diagnosis, the p value was calculated from the chi-square goodness of fit test.

|  | Timepoint | ADNI value | AIBL value | P value |
| --- | --- | --- | --- | --- |
| AGE | 1 | 74.4±9.8 | 74.5±9.8 | 0.99 |
|  | 2 | 76.1±9.8 | 76.1±9.9 | 0.99 |
| MMSE | 1 | 27.9±2.8 | 27.9±2.8 | 1.00 |
|  | 2 | 27.8±2.8 | 27.8±2.8 | 1.00 |
| AD diagnosis | 1 | 57%-43%-0% | 57%-43%-0% | 1.00 |
|  | 2 | 57%-43%-0% | 57%-43%-0% | 1.00 |
| Sex | - | 88% | 88% | 1.00 |

**Table S6.** ADNI-AIBL matching results for participants having 1 time point (scan). For clinical diagnosis in the table, the percentage is showed as CN%-MCI%-AD%. For sex in the table, the portion is the ratio of male subjects. For Age/MMSE, the p value was calculated from a two-sample t-test. For Sex/AD diagnosis, the p value was calculated from the chi-square goodness of fit test.

|  | Timepoint | ADNI value | AIBL value | P value |
| --- | --- | --- | --- | --- |
| AGE | 1 | 74.8±5.9 | 74.8±5.9 | 1.00 |
| MMSE | 1 | 27.3±3.9 | 27.3±3.9 | 0.98 |
| AD diagnosis | 1 | 68%-19%-13% | 68%-19%-13% | 1.00 |
| Sex | - | 43% | 43% | 1.00 |

**Table S7.** ADNI-MACC matching results for participants having 3 time points (scans). For clinical diagnosis in the table, the percentage is showed as CN%-MCI%-AD%. For sex in the table, the portion is the ratio of male subjects. For Age/MMSE, the p value was calculated from a two-sample t-test. For Sex/AD diagnosis, the p value was calculated from the chi-square goodness of fit test.

|  | Timepoint | ADNI value | MACC value | P value |
| --- | --- | --- | --- | --- |
| **AGE** | 1 | 71.5±6.8 | 72.3±6.7 | 0.67 |
|  | 2 | 73.5±6.8 | 73.8±6.8 | 0.91 |
|  | 3 | 75.9±6.9 | 75.5±6.6 | 0.81 |
| **MMSE** | 1 | 26.9±3.7 | 27.0±3.5 | 0.94 |
|  | 2 | 26.1±4.5 | 26.1±4.5 | 0.98 |
|  | 3 | 24.9±6.3 | 25.2±6.3 | 0.87 |
| **AD diagnosis** | 1 | 39%-46%-15% | 36%-54%-10% | 0.72 |
|  | 2 | 43%-36%-21% | 46%-36%-18% | 0.88 |
|  | 3 | 43%-36%-21% | 46%-32%-22% | 0.91 |
| **Sex** | - | 57% | 57% | 1.00 |

**Table S8.** ADNI-MACC matching results for participants having 2 time points (scans). For clinical diagnosis in the table, the percentage is showed as CN%-MCI%-AD%. For sex in the table, the portion is the ratio of male subjects. For Age/MMSE, the p value was calculated from a two-sample t-test. For Sex/AD diagnosis, the p value was calculated from the chi-square goodness of fit test

|  | Timepoint | ADNI value | MACC value | P value |
| --- | --- | --- | --- | --- |
| **AGE** | 1 | 73.6±5.7 | 73.9±5.6 | 0.78 |
|  | 2 | 75.8±5.6 | 75.5±5.6 | 0.71 |
| **MMSE** | 1 | 24.7±4.9 | 24.8±4.6 | 0.86 |
|  | 2 | 23.4±6.9 | 23.5±6.6 | 0.91 |
| **AD diagnosis** | 1 | 35%-38%-27% | 35%-40%-25% | 0.80 |
|  | 2 | 37%-30%-33% | 37%-35%-28% | 0.49 |
| **Sex** | - | 51% | 58% | 0.20 |

**Table S9.** ADNI-MACC matching results for participants having 1 time points (scans). For clinical diagnosis in the table, the percentage is showed as CN%-MCI%-AD%. For sex in the table, the portion is the ratio of male subjects. For Age/MMSE, the p value was calculated from a two-sample t-test. For Sex/AD diagnosis, the p value was calculated from the chi-square goodness of fit test.

|  | Timepoint | ADNI value | MACC value | P value |
| --- | --- | --- | --- | --- |
| AGE | 1 | 75.7±6.7 | 75.7±6.7 | 0.97 |
| MMSE | 1 | 21.0±5.9 | 21.0±5.9 | 0.94 |
| AD diagnosis | 1 | 14%-34%-52% | 14%-38%-48% | 0.64 |
| Sex | - | 52% | 56% | 0.34 |

**Table S10.** Comparison between ComBat with 4 covariates and other harmonization models for ADNI-AIBL. The first row is dataset prediction accuracy, the second row is clinical diagnosis prediction accuracy, and the last row is MMSE prediction mean absolute error (MAE). Within each row, the p values correspond to the difference between ComBat with four covariates and the other models. P values significant after FDR correction (q < 0.05) are bolded.

| ADNI-AIBL | ComBat<br>(AGE+SEX+MMSE+DX) | Models to Compare |  |  |
| --- | --- | --- | --- | --- |
|  |  | ComBat<br>(AGE+SEX) | cVAE | gcVAE |
| Dataset Pred Acc<br>(Mean±Std) | 0.70±0.38 | 0.62±0.41 | 0.60±0.38 | 0.60±0.38 |
|  | p values | **1e-4** | **1e-4** | **1e-4** |
| Clinical Diagnosis Pred<br>Acc (Mean±Std) | 0.45±0.33 | 0.41±0.34 | 0.26±0.29 | 0.69±0.41 |
|  | p values | **0.0014** | **1e-4** | **1e-4** |
| MMSE Pred MAE<br>(Mean±Std) | 1.82±2.48 | 1.82±2.07 | 2.09±3.29 | 1.97±2.93 |
|  | p values | 0.9582 | **0.0013** | **0.0290** |

**Table S11.** Comparison between ComBat with 4 covariates and other harmonization models for ADNI-MACC. The first row is dataset prediction accuracy, the second row is clinical diagnosis prediction accuracy, and the last row is MMSE prediction mean absolute error (MAE). Within each row, the p values correspond to the difference between ComBat with four covariates and the other models. P values significant after FDR correction (q < 0.05) are bolded.

| ADNI-MACC | ComBat<br>(AGE+SEX+MMSE+DX) | Models to Compare |  |  |
| --- | --- | --- | --- | --- |
|  |  | ComBat<br>(AGE+SEX) | cVAE | gcVAE |
| Dataset Pred Acc<br>(Mean±Std) | 0.75±0.36 | 0.72±0.39 | 0.60±0.39 | 0.60±0.40 |
|  | p values | **0.0086** | **1e-4** | **1e-4** |
| Clinical Diagnosis Pred<br>Acc (Mean±Std) | 0.49±0.35 | 0.42±0.35 | 0.36±0.26 | 0.49±0.30 |
|  | p values | **1e-4** | **1e-4** | 0.7204 |
| MMSE Pred MAE<br>(Mean±Std) | 4.95±4.57 | 5.61±5.03 | 6.01±5.50 | 4.25±3.57 |
|  | p values | **1e-4** | **1e-4** | **1e-4** |

**Table S12.** Dataset prediction accuracies with p values of differences between different approaches for unmatched ADNI and AIBL participants. Statistically significant p values after FDR (q < 0.05) corrections are bolded. This is the same as Table 3, except that the roles of matched and unmatched participants were swapped.

| Dataset Prediction Accuracies<br>(mean ± std) | p values |  |  |  |
| --- | --- | --- | --- | --- |
|  | Unharm | ComBat | cVAE | gcVAE |
| Unharmonized (0.991±0.066) |  | **1e-4** | **1e-4** | **1e-4** |
| ComBat (0.676±0.293) |  |  | **1e-4** | **1e-4** |
| cVAE (0.467±0.174) |  |  |  | **1e-4** |
| gcVAE (0.514±0.175) |  |  |  |  |

**Table S13.** Dataset prediction accuracies with p values of differences between different approaches for unmatched ADNI and MACC participants. Statistically significant p values after FDR (q < 0.05) corrections are bolded. This is the same as Table 4, except that the roles of matched and unmatched participants were swapped.

| Dataset Prediction Accuracies<br>(mean ± std) | p values | | | |
| --- | --- | --- | --- | --- |
|  | Unharm | ComBat | cVAE | gcVAE |
| Unharmonized (0.983±0.083) | | **1e-4** | **1e-4** | **1e-4** |
| ComBat (0.720±0.284) | | | **1e-4** | **1e-4** |
| cVAE (0.412±0.139) | | | | **1e-4** |
| gcVAE (0.466±0.176) | | | | |

**Table S14.**
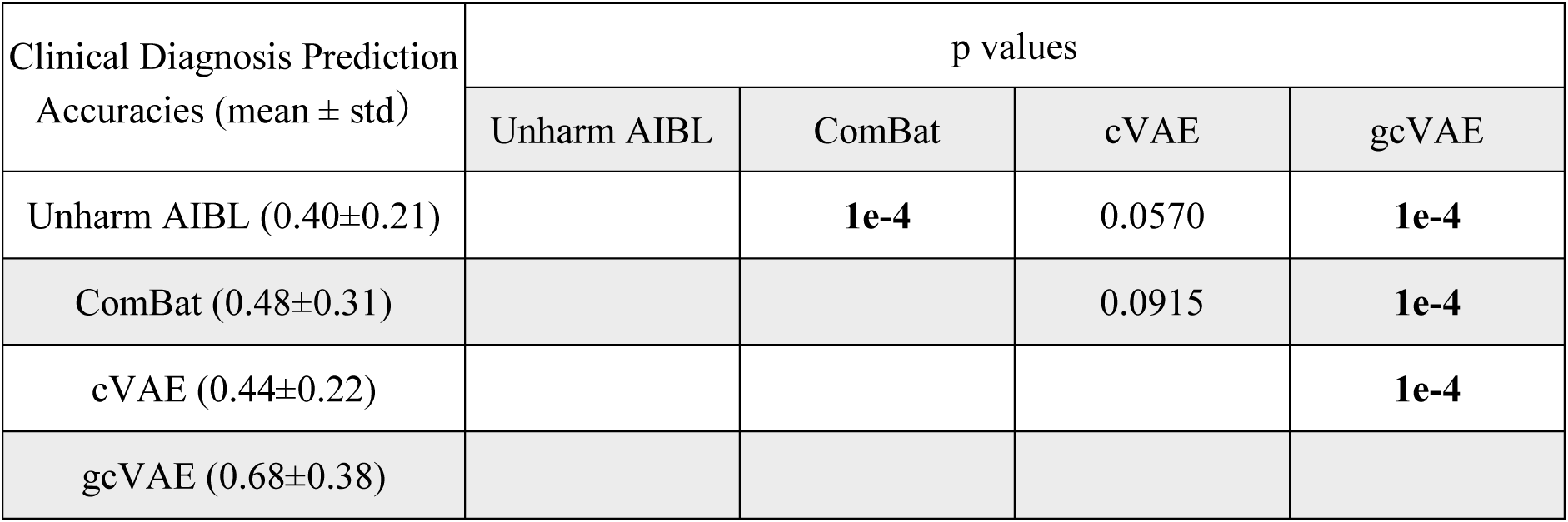
Clinical diagnosis prediction accuracies with p values of differences between different approaches for unmatched AIBL participants. Statistically significant p values after FDR (q < 0.05) corrections are bolded. This is the same as Table 5, except that the roles of matched and unmatched participants were swapped.

**Table S15.** Clinical diagnosis prediction accuracies with p values of differences between different approaches for unmatched MACC participants. Statistically significant p values after FDR (q < 0.05) corrections are bolded. This is the same as Table 6, except that the roles of matched and unmatched participants were swapped.

| Clinical Diagnosis Prediction<br>Accuracies (mean ± std) | p values | | | |
| --- | --- | --- | --- | --- |
|  | Unharm MACC | ComBat | cVAE | gcVAE |
| Unharm MACC (0.47±0.29) | | **0.0021** | 0.2552 | **1e-4** |
| ComBat (0.53±0.31) | | | **2e-4** | 0.1374 |
| cVAE (0.44±0.16) | | | | **1e-4** |
| gcVAE (0.56±0.27) | | | | |

**Table S16.**
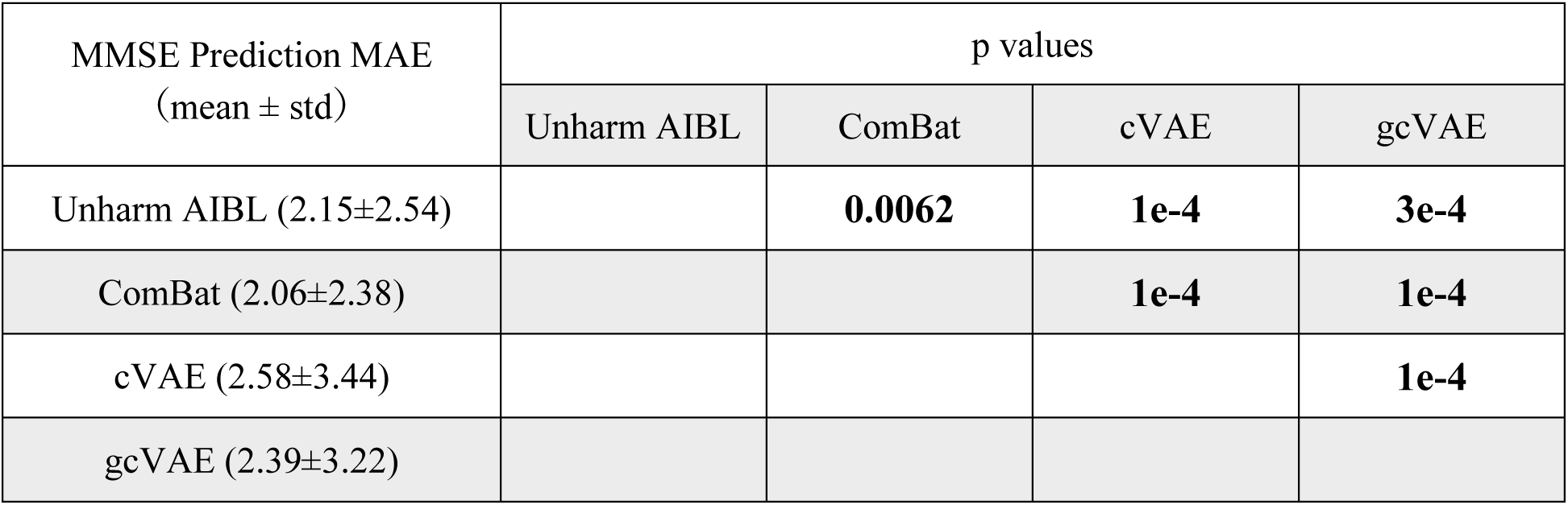
MMSE prediction errors with p values of differences between different approaches for unmatched AIBL participants. Statistically significant p values after FDR (q < 0.05) corrections are bolded. This is the same as Table 7, except that the roles of matched and unmatched participants were swapped.

**Table S17.** MMSE prediction errors with p values of differences between different approaches for unmatched MACC participants. Statistically significant p values after FDR (q < 0.05) corrections are bolded. This is the same as Table 8, except that the roles of matched and unmatched participants were swapped.

| MMSE Prediction MAE<br>(mean ± std) | p values | | | |
| --- | --- | --- | --- | --- |
|  | Unharm MACC | ComBat | cVAE | gcVAE |
| Unharm MACC (6.90±4.49) | | **1e-4** | 0.4411 | **1e-4** |
| ComBat (5.92±4.16) | | | **1e-4** | **1e-4** |
| cVAE (7.00±5.43) | | | | **1e-4** |
| gcVAE (5.03±3.37) | | | | |

**Figure S1.**
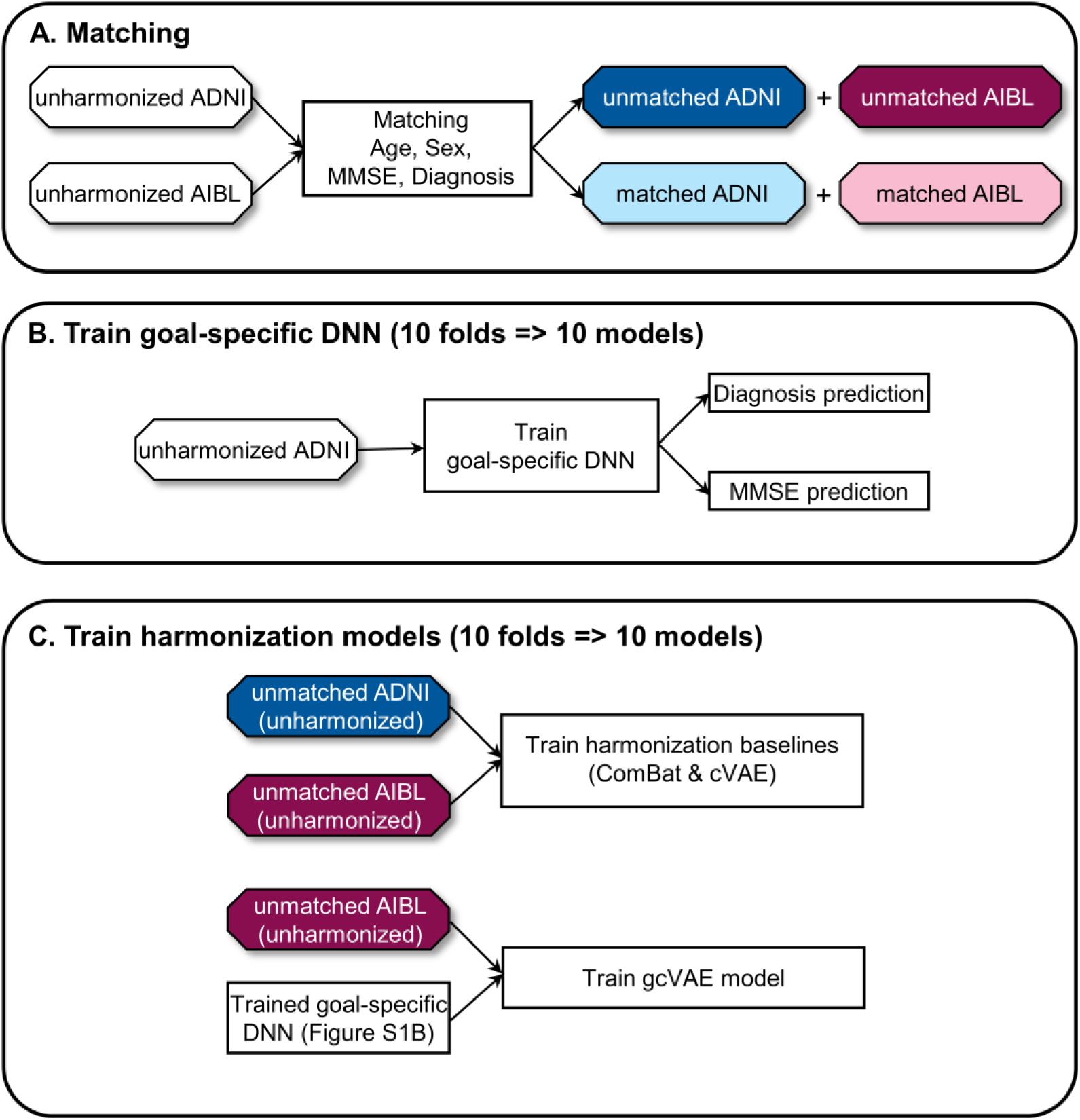
Workflow of control analysis for data matching and model training. We illustrate the workflow using ADNI and AIBL. The same procedure was applied to ADNI and MACC. The workflow is the same as Figure 1 except the role of matched and unmatched participants are swapped in panel C. Furthermore in the case of panel B, all unharmonized data from all ADNI participants was used to train the goal-specific DNN because there are too few matched participants to train the goal-specific DNN.

**Figure S2.**
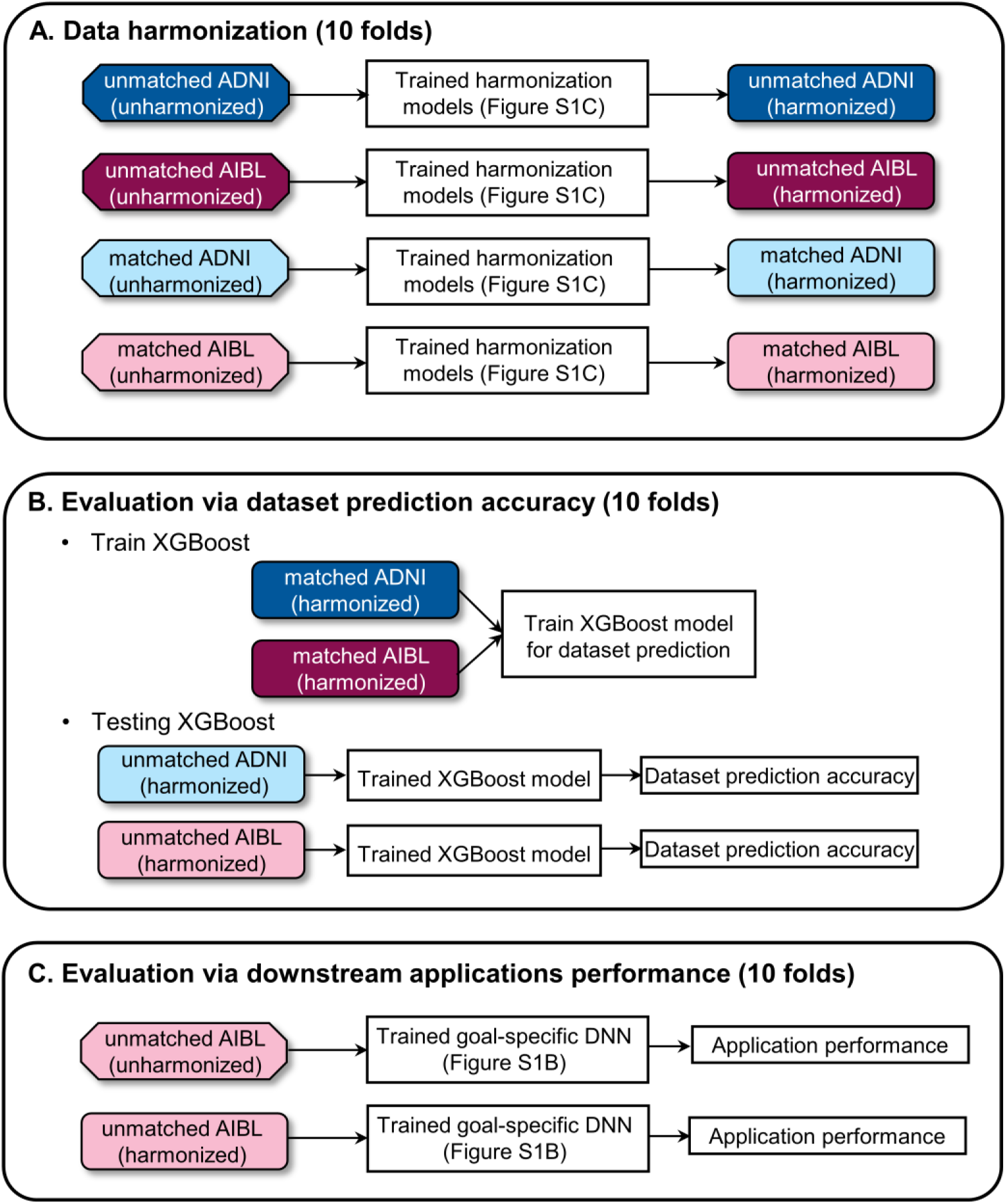
Workflow of control analysis for data harmonization and performance evaluation. We illustrate the workflow using ADNI and AIBL. The same procedure was applied to ADNI and MACC. The workflow is the same as Figure 2 except the role of matched and unmatched participants are swapped in panels B and C. Note that in panel C (compared with Figure 2C), the prediction performance of unmatched unharmonized ADNI and unmatched unharmonized AIBL participants were not comparable, so the downstream application performance was only evaluated on unmatched unharmonized AIBL and unmatched harmonized AIBL data.

**Figure S3.**
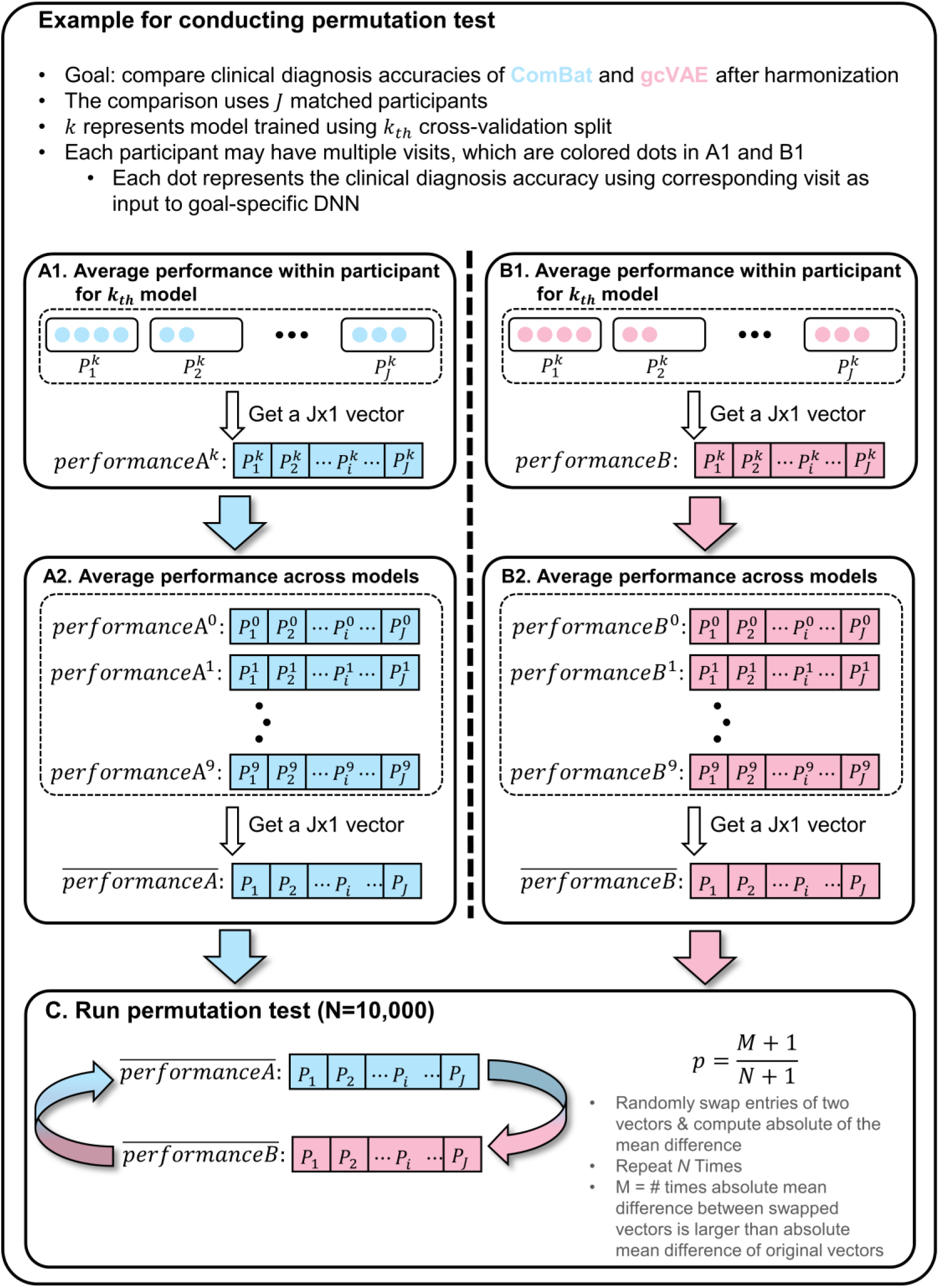
Illustration of permutation test for comparing clinical diagnosis accuracies of ComBat and gcVAE. (A1) For a given model, we averaged the clinical diagnosis accuracies within each participant for ComBat. (B1) Same as A1 but for gcVAE. (A2) Averaging the clinical diagnosis accuracies across the 10 models within each participant. (B1 & B2) Same as A1 and A2 but for gcVAE. (C) Permute 10,000 times to obtain p value.

**Figure S4.**
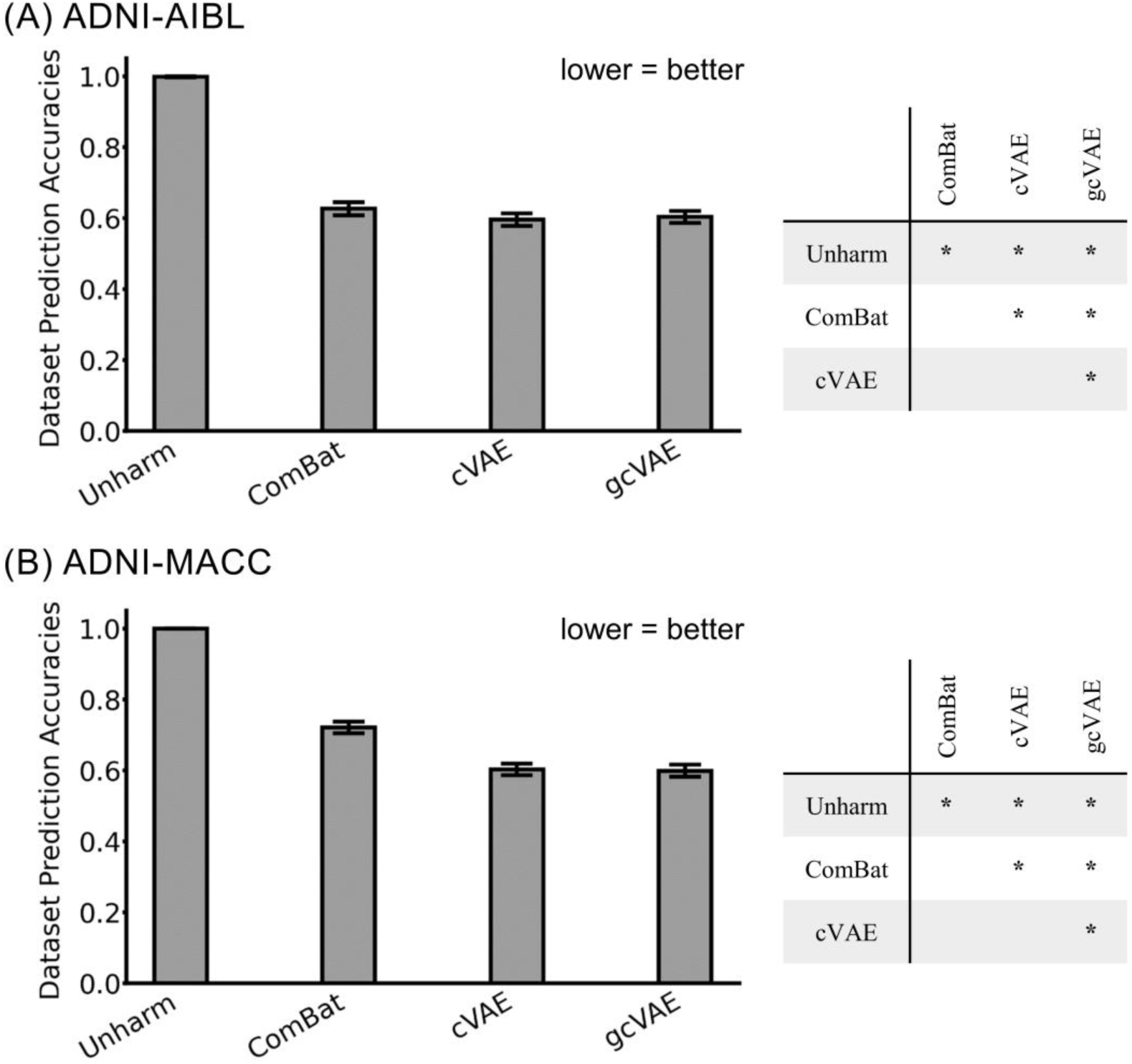
Dataset prediction accuracies when harmonization models were trained on matched data and evaluation was performed on unmatched data. (A) Left: Dataset prediction accuracies for unmatched ADNI and AIBL participants. Right: p values of differences between different approaches. "*" indicates statistical significance after surviving FDR correction (q < 0.05). "n.s." indicates not significant. (B) Same as (A) but for unmatched ADNI and MACC participants. All p values are reported in Tables S12 and S13. This is the same as Figure 5, except that the roles of matched and unmatched participants were swapped.

**Figure S5.**
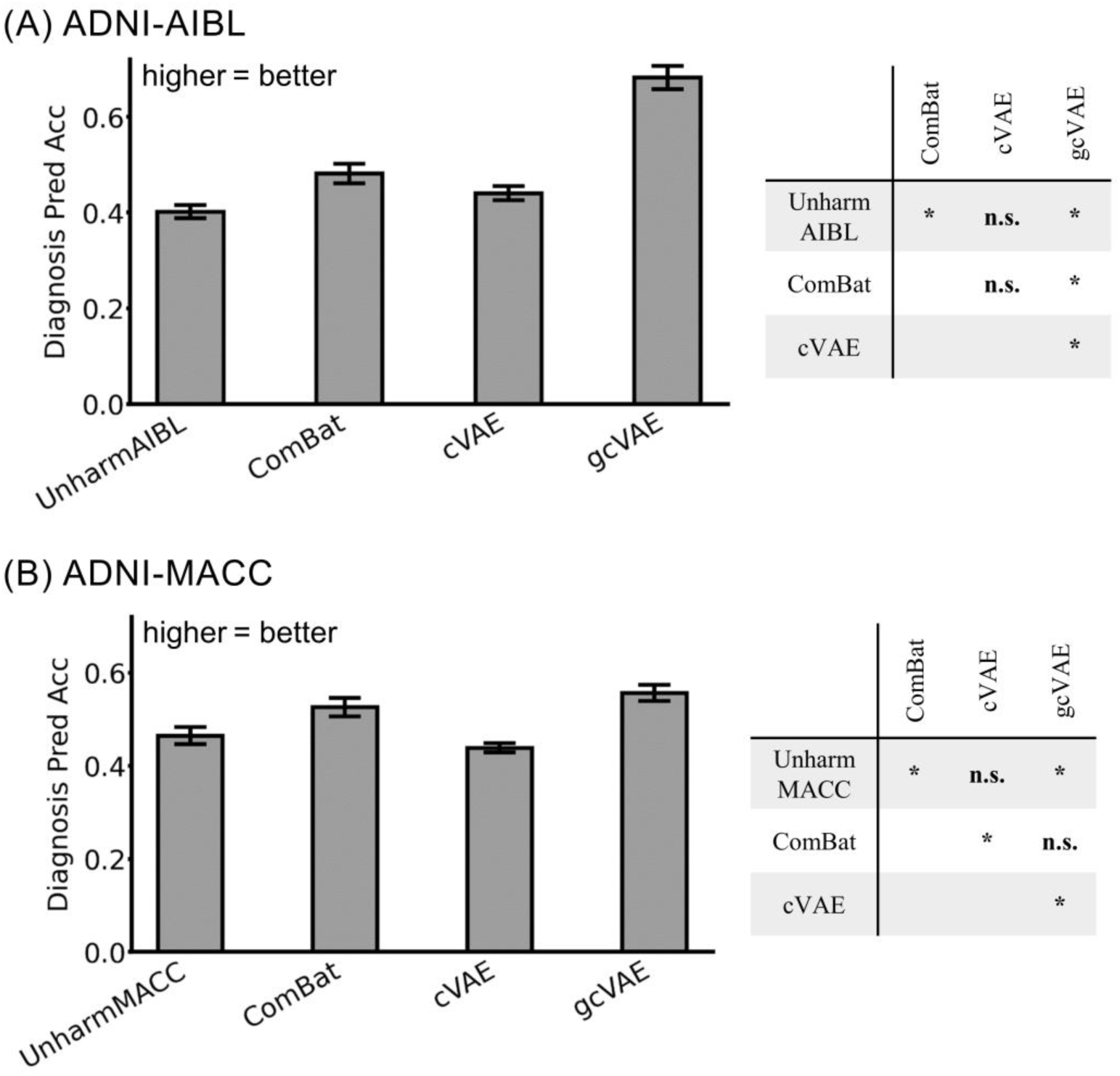
Clinical diagnosis prediction accuracies when harmonization models were trained on matched data and evaluation was performed on unmatched data. (A) Left: Clinical diagnosis prediction accuracies for unmatched AIBL participants. Right: p values of differences between different approaches. "*" indicates statistical significance after surviving FDR correction (q < 0.05). "n.s." indicates not significant. (B) Same as (A) but for unmatched MACC participants. All p values are reported in Tables S14 and S15. This is the same as Figure 6, except that the roles of matched and unmatched participants were swapped.

**Figure S6.**
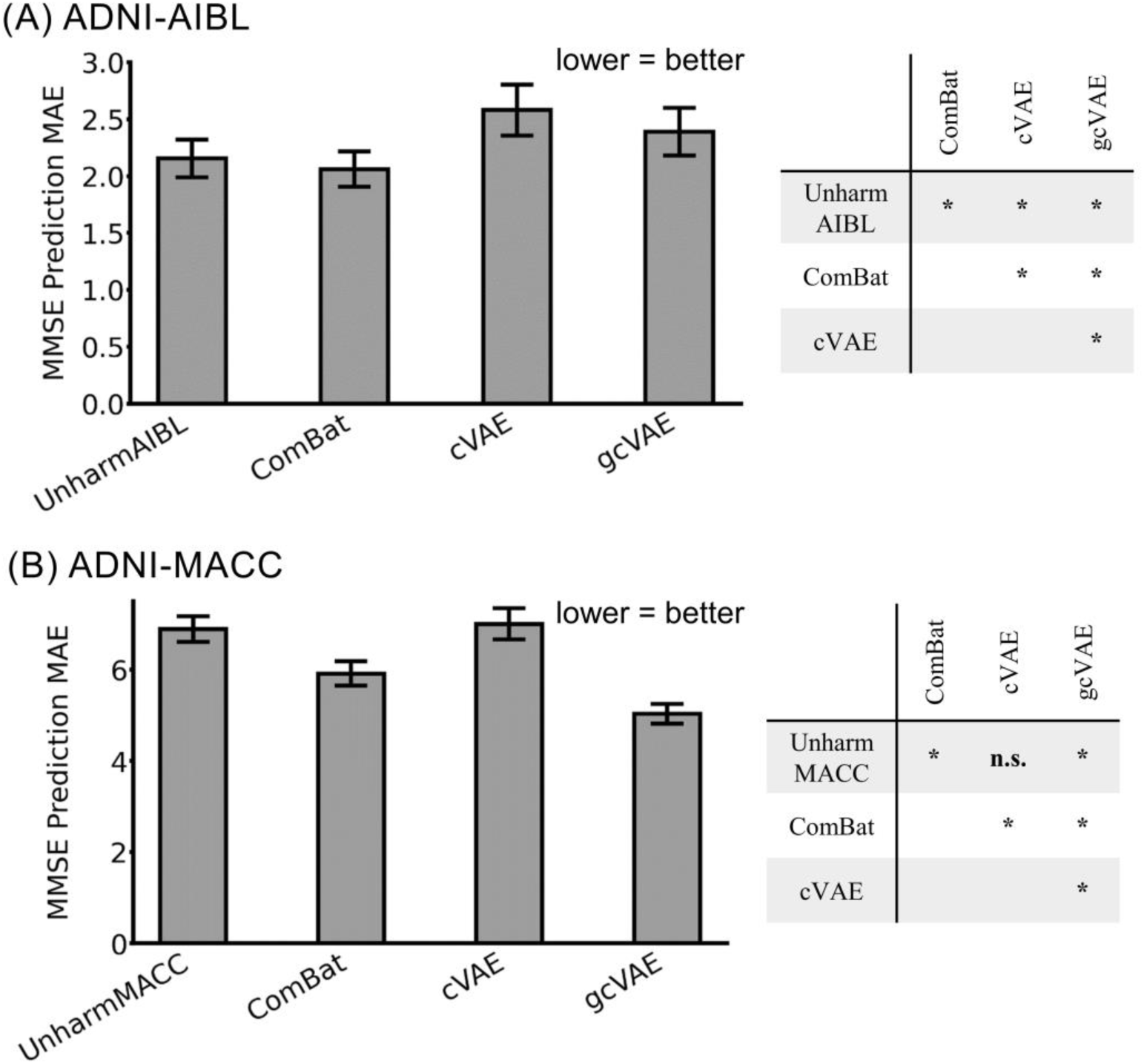
MMSE prediction errors as measured by mean absolute error (MAE) when harmonization models were trained on matched data and evaluation was performed on unmatched data. (A) Left: MMSE prediction errors for unmatched AIBL participants. Right: p values of differences between different approaches. "*" indicates statistical significance after surviving FDR correction (q < 0.05). "n.s." indicates not significant. (B) Same as (A) but for unmatched MACC participants. All p values are reported in Tables S16 and S17. This is the same as Figure 7, except that the roles of matched and unmatched participants were swapped.

